# Targeting proteasome composition reprograms tumor-intrinsic inflammation through selective degradation of STAT1

**DOI:** 10.64898/2026.10.15.759172

**Authors:** Merav D. Shmueli, Yongfeng Tao, Ekampreet Singh, Matthias P Kramer, Anat Abramov, Tuval Aizik, Daoud Sheban, Arseniy Lobov, Karin Goldberg, Angham Ibraheem, Ronen Shteinvil, Aaron Javitt, Yishai Levin, Michael A. Schafroth, Daisuke Ogasawara, Jacob Gross, Frank Adolf, Bruno Melillo, Brenda A Schulman, Benjamin Cravatt, Yifat Merbl

## Abstract

Proteasome regulatory particles determine cellular protein degradation capacities. Yet specific ubiquitin-independent degradation targets of individual proteasome regulatory particles, and physiological roles of such pathways remain largely elusive. Here, through a chemical biology-driven approach, we identify a covalent small molecule that selectively targets a conformationally dynamic pocket in the non-catalytic proteasome regulator PSME4 and modulates tumor cell proteasome targets. Using this chemical probe together with orthogonal genetic perturbations, we show that blocking or reducing PSME4 stabilizes STAT1, restores interferon-associated transcription, enhances antigen presentation, and disrupts IL-6/STAT3 signaling, collectively reprogramming tumor inflammation and suppressing tumor growth *in vivo*. These findings define a PSME4-STAT1-STAT3 axis that couples proteasomal degradation to antigen presentation and cytokine signaling and provide a framework for pharmacologically targeting non-catalytic proteasome regulators to reprogram tumor immunity. More broadly, our findings reveal that proteasome composition can actively govern the balance between immunostimulatory and immunosuppressive signaling programs.

## Introduction

Proteasomes have been broadly implicated in shaping antitumor immunity through roles spanning antigen processing and presentation, inflammatory signaling, and immune cell activation^1–5^. Proteasome function is established by association with regulatory subunits that physically drive a conformational change required for polypeptide access to the protease catalytic chamber. However, beyond intrinsic proteasome activation, the diverse regulatory subunits have distinct structural, biochemical and cellular properties^1,6–10^. Variation in expression of proteasome regulatory subunits^11,12^ is thought to adapt routes of protein degradation to different cellular states and functions. Although the 19S regulatory particle is well understood to mediate ubiquitin- or midnolin-dependent protein degradation^13,14^, few specific targets of other regulatory subunits are known. In particular, if specific proteasome regulators can select a particular substrate for distinct physiological regulation is largely unknown.

A poorly understood proteasome regulator of emerging importance is PSME4 (aka PA200). To date, PSME4 has largely been associated with turnover of acetylated histones during DNA repair and spermatogenesis^16,17^, or acting on a broad repertoire of cellular proteins^18–21^ to shape the degradation landscape and restrict the processing of peptides for MHC-1 presentation. We recently discovered that PSME4 (PA200) is frequently upregulated in Non-Small Cell Lung Cancer (NSCLC), where it alters proteasome activity and MHC-1 antigen presentation, and accordingly associates with lack of response to immunotherapy^15^. However, it remained unclear how cancer-associated alteration of a single proteasome regulatory subunit, namely PSME4, could have such a pronounced effect on anti-tumor immunity. Could a seemingly generic proteasome regulator control the turnover of specific regulatory proteins that govern tumor-intrinsic inflammatory signaling? Addressing this question requires a pharmacological approach that could complement prolonged genetic perturbation and can help distinguish proximal degradation events from secondary adaptive responses. Such an approach could also provide a means to selectively perturb PSME4-containing proteasomes and potentially sensitize tumors to immunotherapy^15^.

Notably, selective pharmacological modulation of regulatory subunits such as PSME4 has remained challenging because of the absence of tractable small-molecule binding pockets characteristic of conventional drug targets. Traditional active site-directed inhibitors, such as bortezomib and carfilzomib, target the catalytic subunits of the 20S core proteasome, resulting in broad proteasome inhibition that is associated with substantial systemic toxicity and limits their clinical utility to disorders in cell types that are hyper-dependent on proteostasis^22,23^. However, selective engagement of non-catalytic proteasome regulators could instead provide a means to perturb specific proteasome functions without globally suppressing proteasomal degradation.

Recent advances in activity-based protein profiling (ABPP) have substantially expanded the ligandable proteome by identifying covalent small molecules that react with cysteines across diverse protein classes^24–28^. However, the relatively high intrinsic reactivity and broad proteome-wide engagement of many electrophilic compounds can limit their utility as selective probes for mechanistic studies. Here, in characterizing by ABPP a library of stereochemically-defined heterocyclic acrylamide (stereoprobe) that span a range of electrophilicity, we discovered a piperidine acrylamide PipAc2-4 that selectively engages cysteine-1001 (C1001) in PSME4 with high stereo- and proteome-wide selectivity. Combined use of PipAc2-4 and genetic depletion revealed that PSME4 regulates the proteasomal degradation of STAT1 in cancer cells and that perturbing PSME4 restored interferon-associated transcription and antigen presentation, as well as disruption of immunosuppressive IL-6/STAT3 signaling. In vivo, PSME4 targeting remodels the tumor immune microenvironment and restricts tumor growth. Together, these findings establish proteasome composition as an endogenous, non-genetic regulator of tumor inflammatory state, extending its function beyond antigen processing, and identify a non-catalytic proteasome regulator as a pharmacologically tractable node for modulating anti-tumor immunity.

## RESULTS

### Library of covalent ligands designed for reduced reactivity as a means to gain selectivity

We previously generated covalent ligandability maps for azetidine acrylamide (AzAc) stereoprobes in human cancer and immune cells by cysteine-directed activity-based protein profiling (ABPP)^26,29,30^, which revealed that these compounds stereoselectively react with cysteines on proteins from structurally and functionally diverse classes. To design a next-generation version probe set with improved features, including decreased intrinsic potential for cross-reactivity and reduced triggering of electrophilic stress responses, we hypothesized that ring size expansion may alleviate the high intrinsic electrophilicity of AzAcs^31^. We prepared full sets of stereochemically defined five-membered (pyrrolidine, PyAc) and six-membered (piperidine, PipAc) heterocyclic acrylamide stereoprobes displaying appendage groups that matched those found in the AzAc counterparts (**Fig. 1A**). For the Pip series, we generated two regioisomeric substitution patterns referred to as PipAc1 and PipAc2. We also produced alkyne-functionalized analogs for each ring-expanded stereoprobe set for use in protein-directed and gel-ABPP experiments^27^ (**Fig. 1A**). Indeed, both PyAc and PipAc stereoprobe sets showed reduced glutathione and global proteomic (gel-ABPP) reactivity compared to their AzAc counterparts, with PipAc probes exhibiting the lowest intrinsic reactivity (**Fig. 1B**, **Fig. S1A**, and **Supplementary Table 1**).

**Figure 1.**
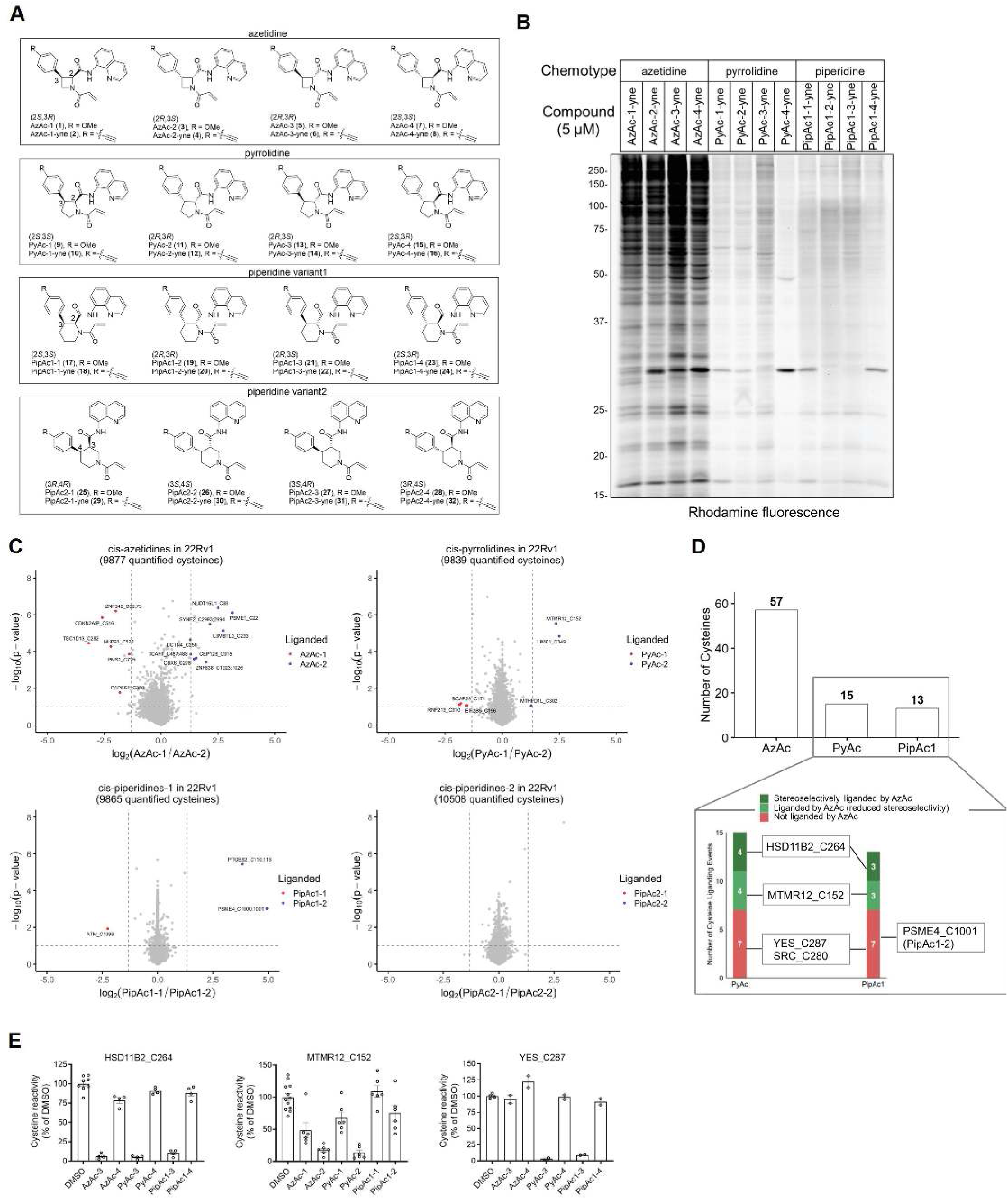
Design and chemical proteomic analysis of ring-expanded electrophilic stereoprobes. (A) Structures of azetidine (four-membered, Az), pyrrolidine (five-membered, Py), and piperidine (six-membered, Pip) acrylamide (Ac) stereoprobes used in this study, including parent compounds and alkynylated analogs. (B) Gel-ABPP data for alkynylated AzAc, PyAc, or PipAc1 stereoprobes (5 μM, 3 h) in HEK293T cells. Stereoprobe-modified proteins were visualized by copper-catalyzed azide alkyne cycloaddition (CuAAC or click) coupling^57,58^ to a rhodamine-azide reporter tag, separation by SDS-PAGE, and visualized by in-gel fluorescence^59^. The data shown are from a single experiment, representative of three independent experiments. (C) Volcano plots of cysteine-directed ABPP data for cis-AzAc, PyAc, PipAc1, and PipAc2 stereoprobes (20 µM, 3 h) in 22Rv1 cells. In brief, ABPP experiments were performed by treating cells with the stereoprobes followed by exposure to a broad-spectrum cysteine-reactive iodoacetamide-desthiobiotin (IA-DTB) probe, such that a stereoselective reduction in IA-DTB labeling identifies stereoprobe-liganded cysteines. Cysteines showing ≥66.7% signal reduction versus DMSO and ≥2.5-fold stereoselectivity between enantiomeric pairs are highlighted in red/blue (two-tailed Student’s t test). Data represents average values from 4-6 independent experiments. (D) (Top) Total number of stereoselectively liganded cysteines for each ring-size chemotype (AzAc, PyAc, PipAc1). (Bottom) Distribution of stereoselective liganding events for PyAc and PipAc1 categorized by AzAc liganding status. Data represents averages from 4-6 independent experiments. (E) Representative examples of stereoselective liganding events from different categories: HSD11B2_C264, liganded by all three ring-size probes; MTMR12_C152, liganded preferentially by ring-expanded stereoprobe PyAc relative to AzAc; and YES1_C287, liganded exclusively by ring-expanded stereoprobes PyAc and PipAc1. Data are from 22Rv1 cells (average ± SEM, n = 2-6).

Evaluating the >20,000 cysteines selectively modified by the ring-expanded stereoprobe sets during cysteine-directed ABPP in two human cancer cell lines revealed both that the ring size expanded PyAc or PipAc stereoprobes produced substantially fewer liganding events than the AzAc stereoprobes (15 and 13, versus 57, respectively; **Fig. 1C-D**, **Fig. S1B**, and **Supplementary Dataset 1**). Moreover, ring expansion did not simply reduce the proteome-wide engagement of stereoprobes: a subset of cysteines showed preferential liganding by the less reactive PyAc or PipAc probes (**Fig. 1D-E**). These interactions consequently exhibited improved proteome-wide selectivity (**Fig. 1C, D** and **Fig. S1B**).

### Selective chemical targeting of PSME4

Among the proteins preferentially liganded by the PipAc stereoprobes was the proteasome regulator PSME4. Mass spectrometry revealed that PSME4 was stereoselectively and site-specifically engaged on a peptide containing two cysteines, C1000 and C1001, by individual stereoprobes from each set of PipAc regioisomers (PipAc1-2 and PipAc2-4; **Fig. 2A, B**). We verified these stereoprobe liganded profiles with recombinant PSME4 expressed in HEK293T cells by gel-ABPP using alkynylated PipAc stereoprobes (**Fig. 2C** and **Fig. S2A**). Site-directed mutagenesis revealed that PipAc stereoprobe reactivity was lost for the C1001A-PSME4, but not the C1000A-PSME4 mutant (**Fig. 2C**). We therefore conclude that the PipAc stereoprobes react with C1001 of PSME4, which is a residue conserved across vertebrates (**Fig. 2C, S2B**).

**Figure 2.**
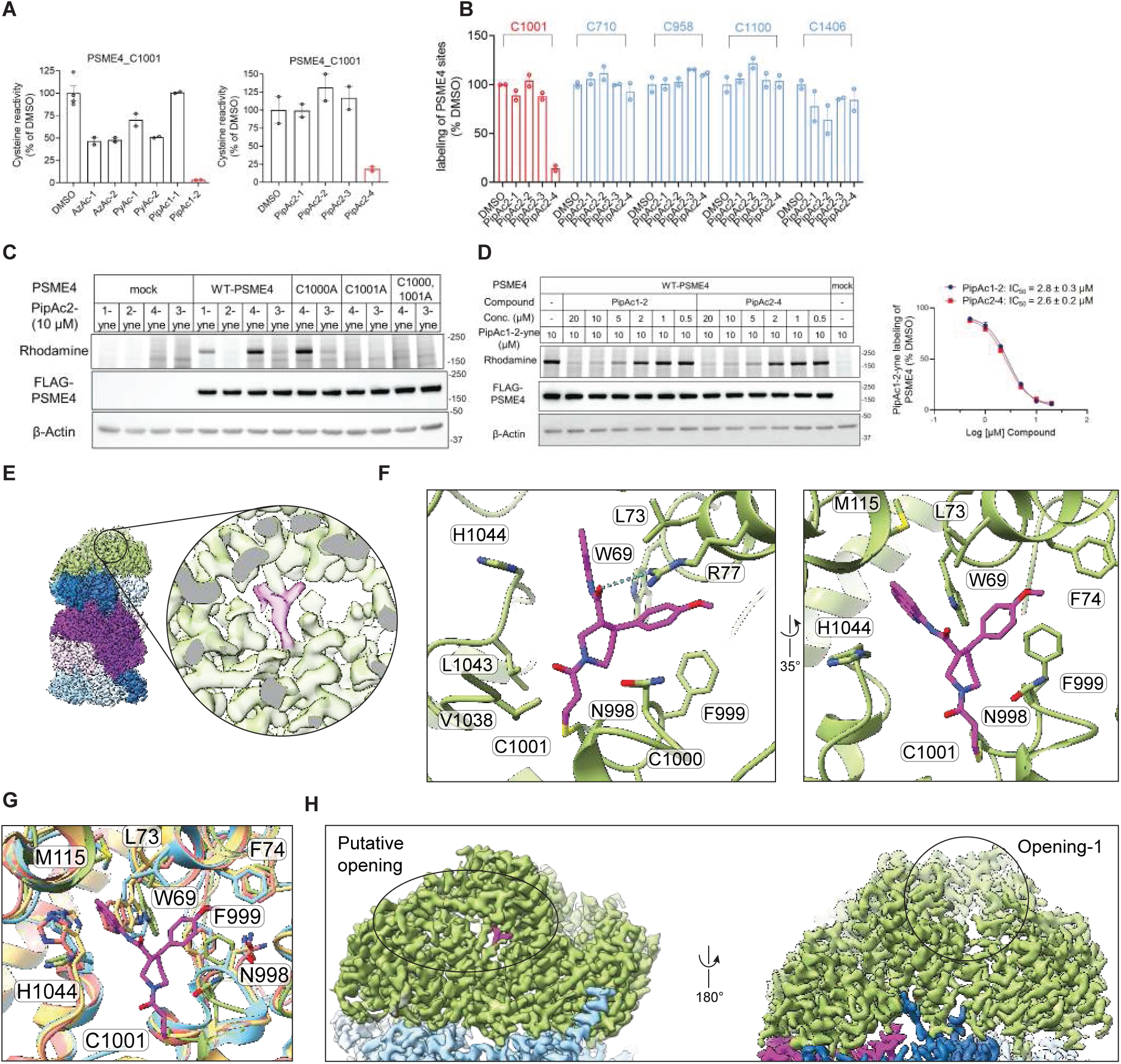
Stereoselective and site-specific liganding of PSME4_C1001 by PipAc stereoprobes. (A) Cysteine-directed ABPP data showing stereoselective liganding of PSME4_C1001 by PipAc1-2 (left) and PipAc2-4 (right) in 22Rv1 cells. Data are average values ± SEM for two to four independent experiments. (B) Cysteine-directed ABPP data for PSME4 from Ramos cells treated with PipAc2 stereoprobes. C1001 (red) and other quantified cysteines (blue) are shown (average ± SEM, n = 2-4). (C) Gel-ABPP of HEK293T cells expressing Flag-tagged WT-PSME4 or the indicated cysteine mutants (C1000A, C1001A, C1000A/C1001A) treated with PipAc2-4-yne (10 μM, 3 h). Data are from a single experiment representative of three independent experiments. (D) Left: Gel-ABPP data showing concentration-dependent engagement of WT-PSME4 by PipAc stereoprobes. WT-PSME4-expressing HEK293T cells were pre-treated with PipAc1-2 and PipAc2-4 (0.5-20 μM, 3 h), followed by PipAc1-2-yne (10 μM, 1 h) and gel-ABPP analysis. Right: Quantification of gel-ABPP providing IC₅₀ curves for PipAc1-2 and PipAc2-4 engagement of WT-PSME4 (mean ± SEM, n = 3). (E) Overview of the cryo-EM map of the PipAc2-4-bound PSME4-20S CP complex. The close-up view highlights the cryo-EM density in a local refined map for the Y-shaped pocket occupied by PipAc2-4 (magenta) in PSME4 (green). (F) Molecular details of the PSME4 pocket bound to PipAc2-4, highlighting the residues of PSME4 that encase PipAc2-4. Hydrogen bonds are shown as dashed lines (cyan). (G) Superimposition of PipAc2-4-bound pocket with all three previously determined models: 6REY (salmon), 6KWY (yellow), and 7NAQ (blue). (H) Cryo-EM map of the PipAc2-4-bound PSME4-20S CP complex highlighting putative openings in PSME4.

Using competitive gel-ABPP assays, we determined IC₅₀ values of ∼2-3 μM for PipAc1-2 and PipAc2-4 engagement of PSME4, whereas other PipAc stereoisomers, as well as azetidine (AzAc-2) and pyrrolidine (PyAc-2), counterparts, showed negligible activity at concentrations up to 10-20 μM (**Fig. 2D** and **Fig. S2C, D**). Revisiting the global reactivity profiles of PipAc1-2 and PipAc2-4, we noted that PipAc2-4 exhibited a superior selectivity profile (**Fig. 1D** and **Fig. S1B**), specifically avoiding cross-reactivity with the prostaglandin synthesis enzyme PTGES2 (**Fig. S2E**). We therefore proceeded with PipAc2-4 for subsequent studies of PSME4.

The PipAc2-4 mode binding to the PSME4-20S CP was revealed by cryo-EM (**Fig. S3** and **Supplementary Table S2**). A focus-refined map over PSME4 at 2.8 Å resolution confirmed covalent binding at C1001, with PipAc2-4 occupying a Y-shaped pocket that accommodates the central piperidine of the small molecule core along with its quinoline and anisole appendages projected in opposing directions (**Fig. 2E, F**). The distinct PSME4 features on the two sides of the “Y” explain both the targeting selectivity of PipAc2-4, and the inactivity of the enantiomer PipAc2-3. On one side, the quinoline is encased by W69, L73, M115 and H1044, and, on the other, the anisole is embraced by aromatic residues W69, F74 and F999 and engages in aromatic stacking with R77 and further hydrophobic interactions with L73 (**Fig. 2E, F**). Interestingly, many of the side chains interacting with PipAc2-4 are substantially rearranged compared to previous PSME4-20S CP structures^8,32,33^ (7NAQ, 6KWY and 6REY), which are all notably similar to each other in this region (**Fig. 2G** and **S4A-C**).

Beyond the local conformational changes in the PipAc2-4-binding site, the remainder of the PSME4-20S CP complex is largely similar to prior structures^8,32,33^. However, we noted that a region previously regarded as a putative opening^32^, and that is in close proximity to PipAc2-4, displayed heterogeneity in our maps and some differences from previous structures^8,32,33^ (**Fig. 2H** and **S4D**). Also, at the opening 1 portion of PSME4, some features that are well-defined in a subset of prior maps^8,32,33^ are either invisible or only observed as weak density in the PipAc2-4-bound complex (**Fig. S4E**). Since these regions of PSME4 have been proposed as potential substrate entry and/or product exit sites, we considered that the apparent alterations caused by PipAc2-4 binding might impact the substrate selectivity and proteasome activity of the PSME4-20S CP complex.

### Small molecule targeting of PSME4 alters proteasome activity and restores antigen presentation

Having established selective engagement of PSME4, we asked whether PipAc2-4 recapitulates the functional consequences of PSME4 loss. To directly assess catalytic effects, we reconstituted purified 20S-PSME4 complexes *in vitro* (**Fig. S5A-B**). PSME4 increased cleavage of the β1-associated nLPnLD-AMC substrate relative to recombinant 20S proteasome alone (**Fig. 3A-B**). PipAc2-4 reduced this activity in a concentration-dependent manner, whereas related stereoisomers had minimal effects (**Fig. 3C-D** and **Fig. S5C-E**). PipAc2-4 also reduced β2- and β5-associated activities in purified 20S-PSME4 complexes (**Fig. S5F-G**), consistent with attenuation of PSME4-capped proteasome activity. In proteasomes isolated from treated cells, however, PipAc2-4 selectively decreased caspase-like (β1-associated) activity and increased chymotryptic activity, without significantly affecting tryptic-like activity (**Fig. 3E-F** and **Fig. S5H-K**). Thus, covalent PSME4 liganding by PipAc2-4 reshaped proteasome activity in cells rather than causing global catalytic inhibition.

**Figure 3.**
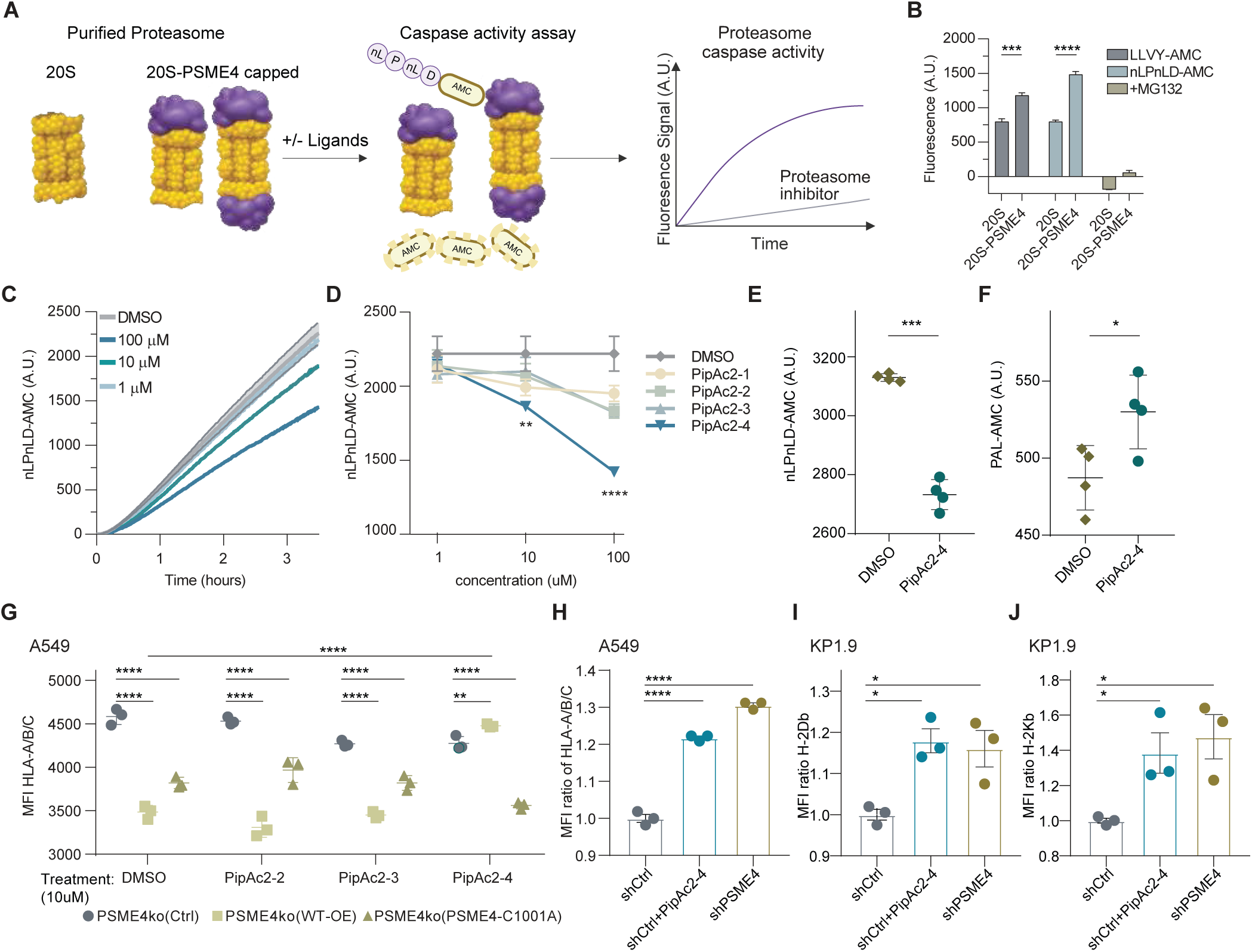
Covalent engagement of PSME4 C1001 alters proteasome activity and increases MHC-I presentation. (A) Schematic of the *in vitro* proteasome activity assay of purified 20S-PSME4 complexes with fluorogenic peptide substrates in the presence or absence of stereoprobes. (B) Cleavage of LLVY-AMC and nLPnLD-AMC by recombinant 20S (Enzo) or purified 20S-PSME4 complexes; MG132 was included as a catalytic proteasome inhibitor control. (C) Time course of nLPnLD-AMC cleavage by purified 20S-PSME4 complexes in the presence of increasing concentrations of PipAc2-4 (mean ± SEM, n = 2). (D) Dose-response analysis of PipAc2-4-dependent suppression of nLPnLD-AMC cleavage (2-way ANOVA with Uncorrected Fisher’s LSD, mean ± SEM, n = 2). (E, F) Proteasomes isolated by PSMA1 immunoprecipitation from A549 cells treated with PipAc2-4 or vehicle (DMSO) and assayed for cleavage of nLPnLD-AMC (E) and PAL-AMC (F) (mean ± SEM, n = 4, two-sided t-test). (G) Surface HLA-A/B/C abundance in PSME4-knockout A549 cells reconstituted with luciferase control, wild-type PSME4, or PSME4 C1001A and treated with DMSO or the indicated PipAc2 stereoisomers (10 μM, 18 h; mean ± SEM, n = 3; two-way ANOVA with Holm-Sidak multiple-comparisons test). (H-J) Surface MHC-I abundance in parental A549 cells (H) and KP1.9 cells (I, J) following PipAc2-4 treatment or PSME4 depletion, as indicated (mean ± SEM, n = 3; one-way ANOVA with Holm-Sidak multiple comparisons test).

We tested if treatment with PipAc2-4 recapitulates the effect of genetic PSME4 depletion on antigen presentation^15^, and examined the effects on PSME4-knockout A549 cells reconstituted with wild-type or mutant PSME4. PipAc2-4 increased surface HLA-A/B/C, whereas cells expressing the C1001A mutant were refractory to the compound (**Fig. 3G-H** and **Fig. S5L-M**), linking the pharmacological effect to engagement of PSME4 C1001. PipAc2-4 similarly increased MHC-I surface abundance in murine KP1.9 cells (**Fig. 3I-J** and **Fig. S5N**). Together, these findings establish PipAc2-4 as a functional probe of PSME4 and demonstrate that covalent liganding of C1001 alters proteasome activity and restores MHC-I antigen presentation.

### Selective chemical targeting reveals PSME4-capped proteasomes promote STAT1 degradation

The ability to acutely and selectively perturb PSME4-capped proteasomes provided a route to identify specific substrates whose degradation could shape tumor-intrinsic inflammatory signaling, as compound treatment could enable the distinction of proximal degradation events from broader adaptive responses associated with prolonged genetic perturbation. Mass spectrometry analysis of proteasome-cleaved peptides (MAPP^34^) after PipAc2-4 treatment identified 550 proteasome-cleaved peptides, 60 proteins, with reduced abundance relative to control (DMSO) (pVal<0.05; FC>1.5) (**Fig. 4A** and **Fig. S6A-B, Supplementary Table S3**). Complementary profiling following PSME4 overexpression identified 480 differentially represented peptides and 80 proteins relative to GFP control (pVal<0.05; FC>1.5) (**Fig. 4B** and **Fig. S6C-D, Supplementary Table S4**). Comparison of the source proteins of the degraded peptides 60 shared candidate proteins among 296 nonredundant significant proteins represented across the pharmacological and overexpression analyses (16.9%; **Fig. 4C** and **Fig. S6E, Supplementary Table S5**). Despite differences in perturbation direction and duration, this convergence identified a restricted set of proteins whose degradation was sensitive to loss of PSME4 levels or activity.

**Figure 4.**
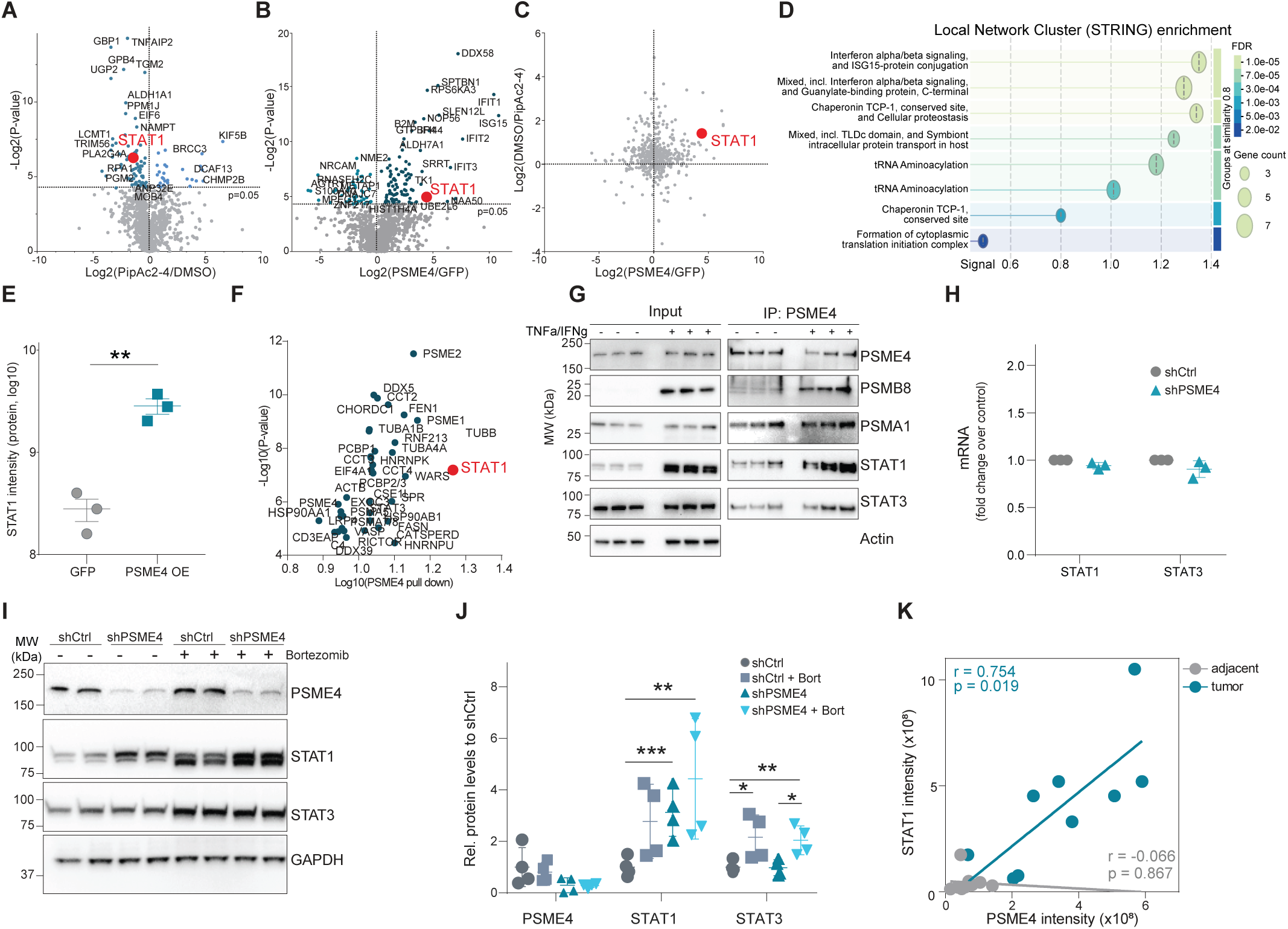
PSME4-capped proteasomes promote STAT1 degradation. (A) MAPP degradome profiling of A549 cells treated with PipAc2-4 for 18 h or DMSO, showing proteasome-cleaved peptides with altered abundance following pharmacological PSME4 perturbation. V-plot presents the protein-level differential between treatments. (B) MAPP profiling of A549 cells overexpressing PSME4 for 48 h relative to GFP control, showing proteasome-cleaved peptides altered by increased PSME4 abundance. V-plot presents the protein-level differential between treatments. (C) Comparison of the pharmacological and PSME4-overexpression datasets identifies convergent candidate proteins, including STAT1. Proteins that were detected in both experiments (a total of 519 proteins) were plotted by Log2 (fold change). (D) Pathway over-representation analysis of differentially degraded proteins following PSME4 overexpression using STRING Local Network Clusters. Bubble size indicates the number of proteins contributing to each pathway, and color denotes the Benjamini-Hochberg-adjusted p value (FDR), revealing enrichment of interferon-associated pathways and interferon-stimulated genes. (E) PSMA1 immunoprecipitation of proteasome complexes from PSME4-overexpressing cells showing increased proteasome-associated STAT1 relative to GFP control (mean ± SEM, n = 3; unpaired two-tailed t test). (F) Mass-spectrometry analysis of immunoprecipitated PSME4 identifies STAT1 together with multiple proteasome subunits among PSME4-associated proteins. (G) PSME4 immunoprecipitation followed by immunoblotting for associated STAT1 under basal conditions and following TNFα/IFNγ stimulation. (H) qPCR analysis of STAT1 and STAT3 transcript abundance in control and PSME4-depleted A549 cells (mean ± SEM, n = 3). (I, J) Immunoblotting (I) and quantification (J) of STAT1 and STAT3 abundance in control and PSME4-depleted A549 cells following bortezomib treatment. Bortezomib-dependent STAT1 accumulation is attenuated following PSME4 depletion (mean ± SEM, n = 4; two-way ANOVA with Tukey’s multiple comparisons test). (K) Correlation between PSME4 abundance and proteasome-associated STAT1 in primary NSCLC tumors and matched adjacent tissue following proteasome immunoprecipitation (n = 9; Pearson correlation coefficients and calculated 2-tailed p values).

STAT1 was selected for further analysis because it was identified across both perturbation datasets and occupies a central role in tumor-intrinsic interferon responses^35^. Loss-of-function alterations in JAK1/2 that impair tumor-cell responsiveness to IFN have been linked to both acquired and primary resistance to PD-1 blockade^36,37^. Conversely, prolonged IFN signaling can promote resistance to immune checkpoint blockade through a STAT1-dependent multigenic program that includes multiple T cell inhibitory ligands^38^. Thus, tumor cells must balance IFN signaling between insufficient and sustained activation, although the endogenous mechanisms that tune the amplitude of this signaling independently of genetic lesions in the pathway remain poorly understood. In particular, whether proteasome composition itself can regulate IFN signaling by controlling STAT1 protein stability is unknown. Yet our pathway analysis of the top differentially degraded proteins following PSME4 overexpression revealed enrichment of interferon α/β signaling and ISG15-associated pathways, including IFIT1, IFIT2, IFI44, and B2M (**Fig. 4D**). Together, these findings implicated PSME4-dependent proteolysis in the regulation of the interferon-responsive state.

Notably, several experiments revealed that STAT1 association with proteasome complexes changed as a function of PSME4 abundance. Immunoprecipitation of endogenous proteasomes through the PSMA1 (α6) subunit showed increased co-association of STAT1 following PSME4 overexpression (**Fig. 4E**). Reciprocal PSME4 immunoprecipitation followed by mass spectrometry identified STAT1 together with multiple proteasome subunits (**Fig. 4F**), and immunoblotting confirmed PSME4-STAT1 co-association under basal conditions and following TNFα/IFNγ stimulation (**Fig. 4G**). STAT1 and STAT3 transcript abundance were unchanged following PSME4 depletion (**Fig. 4H**), supporting post-transcriptional regulation. Consistent with proteasome-dependent degradation, bortezomib increased STAT1 abundance in control cells, whereas this effect was attenuated following PSME4 depletion (**Fig. 4I-J** and **Fig. S6F**). These results support a model in which PSME4-containing proteasomes promote STAT1 turnover.

Finally, to assess relevance in human disease, we analyzed proteasome complexes isolated from primary NSCLC tumors and matched adjacent tissue. Within tumors, PSME4 abundance correlated positively with proteasome-associated STAT1, whereas no correlation was detected in adjacent tissue (Pearson r = 0.754, p = 0.019 versus r = -0.066, p = 0.867; **Fig. 4K**). These data support a tumor-specific association between PSME4 abundance and STAT1 recruitment to proteasome complexes in human NSCLC. Together, these findings identify STAT1 as a PSME4-regulated proteasomal substrate and link altered proteasome composition directly to tumor-intrinsic interferon signaling.

### PSME4-dependent STAT1 degradation shifts tumor inflammatory signaling toward an IL-6/STAT3 state

Having identified STAT1 as a PSME4-regulated substrate, we asked whether its stabilization reprograms downstream inflammatory signaling. Unlike STAT1, total STAT3 abundance remained largely unchanged following PipAc2-4 treatment or genetic PSME4 depletion (**Fig. 5A-B**). We therefore examined whether increased STAT1 abundance altered its association with STAT3. Non-denaturing STAT3 immunoprecipitation showed increased co-association of STAT1 following PSME4 depletion (**Fig. 5C** and **Fig. S7A**), indicating that PSME4 loss alters the composition of STAT-containing signaling complexes.

**Figure 5.**
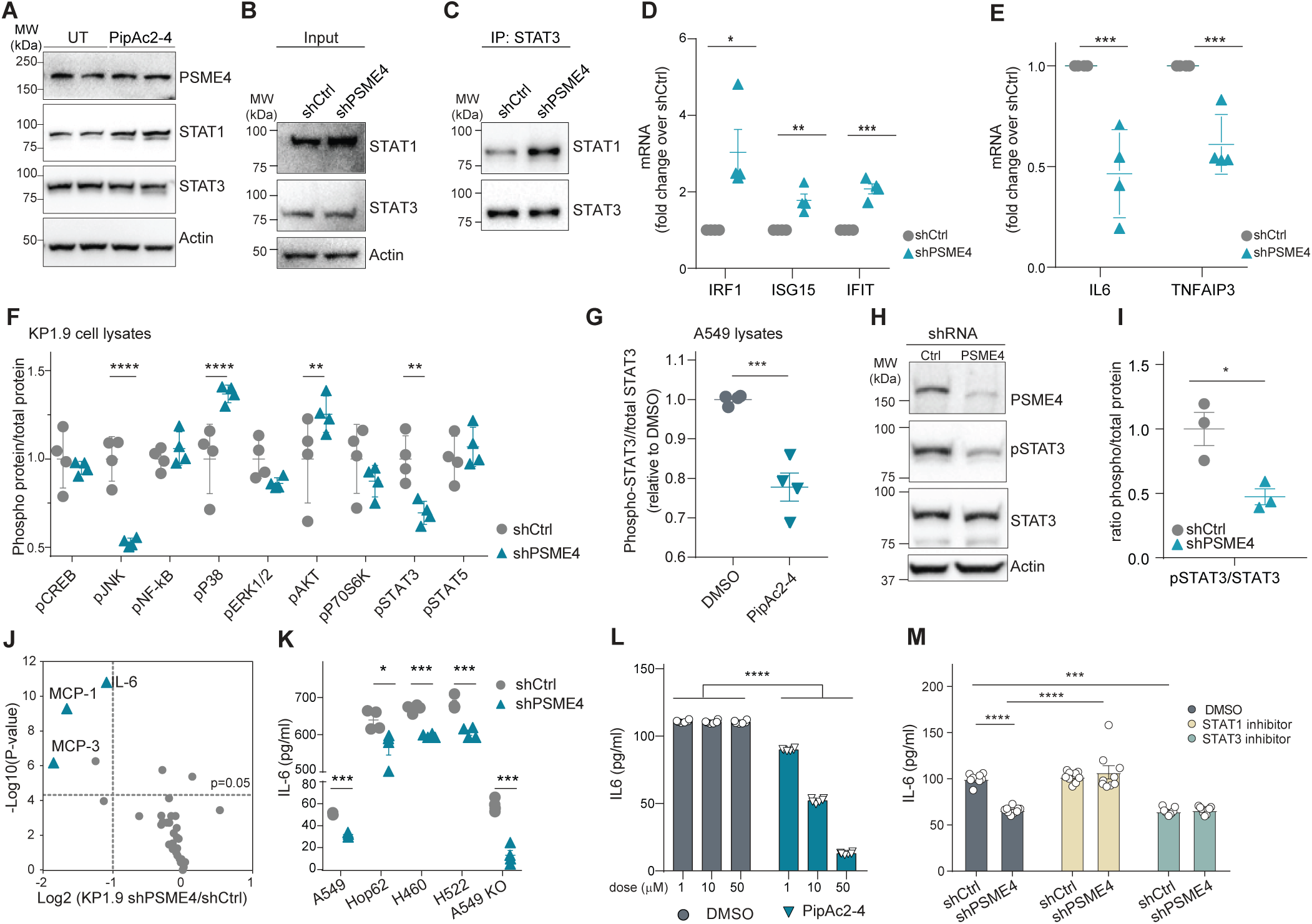
PSME4-dependent STAT1 degradation shifts inflammatory signaling toward an IL-6/STAT3-associated state. (A) Immunoblot analysis of A549 cells treated with PipAc2-4 or DMSO, showing increased STAT1 abundance with largely unchanged total STAT3. (B, C) STAT3 immunoprecipitation under non-denaturing conditions followed by analysis of STAT1 co-association in control and PSME4-depleted cells. (D, E) qPCR analysis showing increased expression of interferon-associated STAT1 targets (IRF1, ISG15, and IFIT1) and reduced expression of TNFAIP3 and IL6 following PSME4 depletion (mean ± SEM, n = 4; unpaired t tests with Holm-Sidak correction for multiple comparisons). (F) Multiplex phospho-signaling analysis compared to total protein abundance following PSME4 depletion, showing changes in phosphorylated STAT3, JNK, and other measured signaling nodes (mean ± SEM, n = 4; multiple unpaired t tests with FDR correction). (G) Multiplex analysis of phosphorylated relative to total STAT3 following PipAc2-4 treatment (mean ± SEM, n = 4; unpaired two-tailed t test). (H, I) Immunoblotting (H) and quantification (I) showing reduced STAT3 phosphorylation relative to total STAT3 following PSME4 depletion, with total STAT3 abundance largely unchanged (mean ± SEM, n = 3; p values as indicated; unpaired two-tailed t test). (J) Multiplex cytokine profiling following PSME4 depletion, with selected differentially abundant cytokines indicated, including reduced IL-6. (K) IL-6 secretion across four human NSCLC cell lines following PSME4 depletion; PSME4-knockout A549 cells are included as an additional genetic perturbation (mean ± SEM, n = 4; p values as indicated; unpaired t tests with Holm-Sidak correction for multiple comparisons). (L) IL-6 ELISA dose-response analysis in A549 cells treated with PipAc2-4 relative to DMSO (mean ± SEM, n = 3; p values as indicated; unpaired t tests with Holm-Sidak correction for multiple comparisons). (M) IL-6 secretion in control and PSME4-depleted A549 cells following treatment with STAT3 inhibitor, fludarabine, or STAT1 inhibitor, nifuroxazide, (mean ± SEM, n = 9; unpaired two-tailed t test).

Consistent with restoration of a STAT1-associated transcriptional program, PSME4 depletion increased expression of IRF1, ISG15, and IFIT1 while reducing IL6 and TNFAIP3 expression (**Fig. 5D-E**). These coordinated changes were consistent with enhanced interferon-associated transcription and reduced STAT3-linked inflammatory signaling. Multiplex phospho-signaling analysis of canonical inflammatory pathways further revealed reduced STAT3 phosphorylation together with additional pathway changes, most prominently reduced JNK phosphorylation, whereas several other measured proteins showed smaller or nonsignificant effects (e.g. pNF-kB, pERK1/2, pSTAT5; **Fig. 5F**). Consistent with the STAT3 result, both PipAc2-4 treatment and genetic PSME4 depletion reduced phosphorylated STAT3 relative to total STAT3 without substantially altering total STAT3 abundance (**Fig. 5G-I** and **Fig. S7B**). Thus, PSME4 loss shifts signaling away from a STAT3-dominant state without globally suppressing inflammatory pathways.

Because persistent STAT3 signaling promotes production of immunosuppressive cytokines, particularly IL-6^39,40^, we examined cytokine secretion following PSME4 depletion. Multiplex cytokine profiling identified IL-6 as one of the most strongly reduced cytokines (**Fig. 5J** and **Fig. S7C**). Following short-term inflammatory stimulation, IL-6 remained significantly reduced, whereas several other measured cytokines increased, including IFN-γ, TNFα, and IL-17A (**Fig. S7D**). Reduced IL-6 secretion following PSME4 depletion was also reproduced across multiple NSCLC human cell lines (**Fig. 5K**). PipAc2-4 also reduced IL-6 secretion in a concentration-dependent manner, whereas related stereoisomers did not (**Fig. 5L** and **Fig. S7E**), linking selective pharmacological engagement of PSME4 to suppression of IL-6 secretion.

Finally, we used pharmacological perturbation of STAT signaling to test whether STAT-associated activity contributes to altered IL-6 secretion following PSME4 depletion. The STAT1-associated inhibitor fludarabine increased IL-6 secretion in PSME4-depleted cells, whereas the STAT3 inhibitor nifuroxazide further reduced IL-6 (**Fig. 5M**). Collectively, these findings support a model in which PSME4-dependent STAT1 turnover attenuates interferon-associated transcription and favors a STAT3/IL-6 inflammatory program, whereas disruption of PSME4 restores STAT1-linked ‘inflammatory competence’.

### Targeting PSME4 remodels tumor inflammation and anti-tumor immunity *in vivo*

We asked whether restoration of tumor-intrinsic interferon competence following PSME4 depletion alters tumor growth, immune composition, and therapeutic response *in vivo*. PSME4-deficient subcutaneous KP1.9 tumors exhibited reduced growth and lower final tumor weights relative to control tumors (**Fig. 6A-B**). Because PSME4 depletion did not measurably alter KP1.9 proliferation under the *in vitro* conditions tested (**Fig. S8A**), the *in vivo* growth difference is unlikely to be explained solely by a constitutive cell-autonomous proliferation defect. Further, as reported in our previous work, the tumor growth attenuation of PSME4-deficient KP1.9 cells was T cell-mediated, in an orthotopic model of NSCLC^15^.

**Figure 6.**
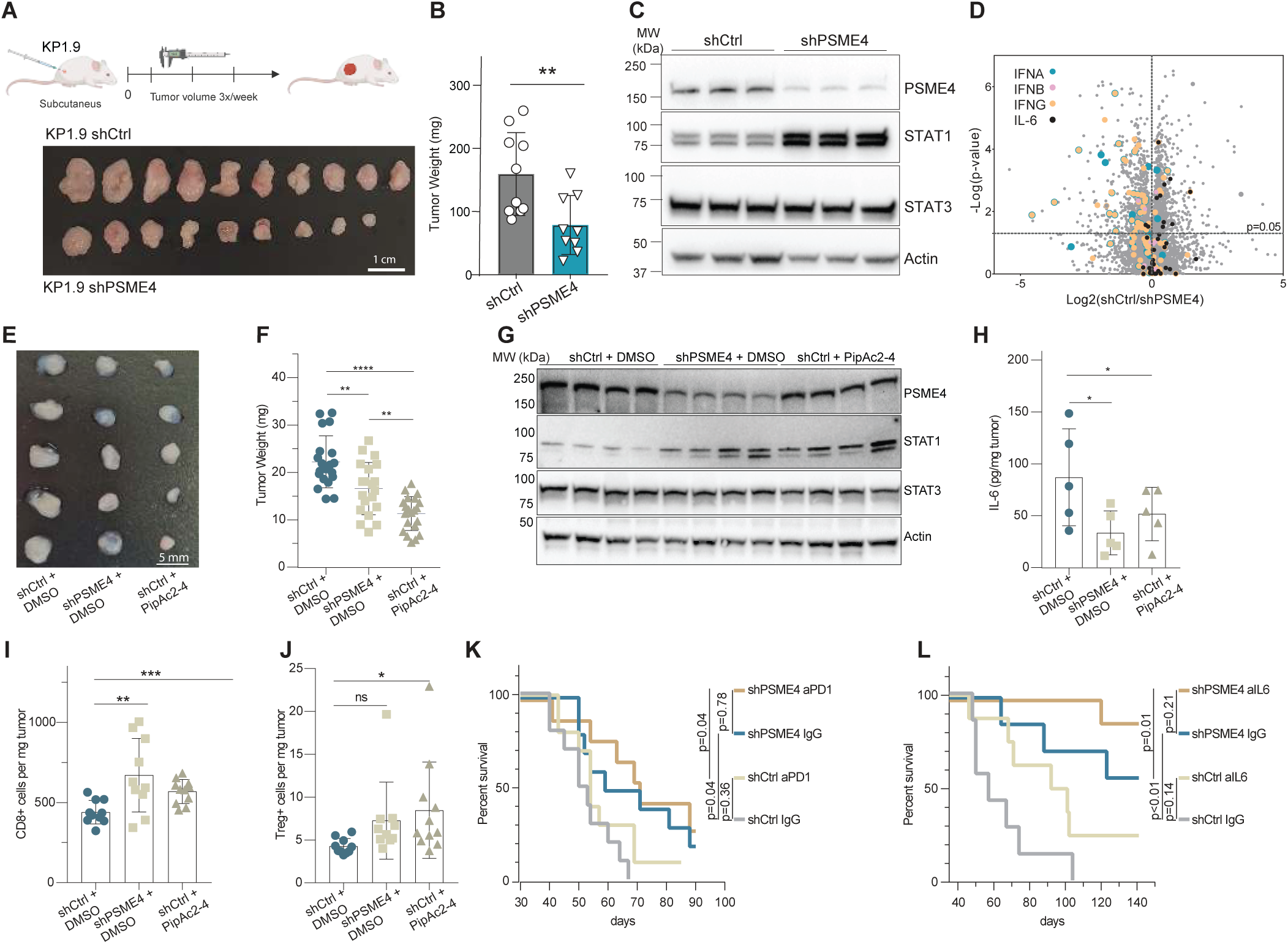
PSME4 depletion remodels tumor inflammation and anti-tumor immunity *in vivo*. (A, B) Analysis of subcutaneous KP1.9 tumors expressing control or PSME4-targeting shRNA. Experimental schematic and representative tumors are shown in (A), and tumor growth is shown in (B) (mean ± SEM; n = 10 for shCtrl and n = 9 for shPSME4; unpaired two-tailed t-test). (C) Immunoblot analysis of resected tumor lysates showing increased STAT1 abundance and largely unchanged total STAT3 following PSME4 depletion. Actin serves as a loading control. (D) Differential proteomic analysis of control and PSME4-depleted tumors, highlighting changes in interferon- and IL-6-associated proteins as indicated by colors. (E, F) Priming-rechallenge model in which mice were initially primed with KP1.9 tumors and, 14 days later, rechallenged with KP1.9 shCtrl cells pretreated with PipAc2-4 or DMSO, or with KP1.9 shPSME4 cells. Tumor mass measured 48 h after rechallenging is shown in (F) (mean ± SEM, n = 20; one-way ANOVA with Tukey’s multiple- comparisons test). (G) Immunoblotting of resected tumors shows STAT1 abundance from the rechallenge model following PipAc2-4 pretreatment or PSME4 depletion. (H) Tumor- associated IL-6 secretion under the indicated conditions (mean ± SEM, n = 5; p values as indicated; one-way ANOVA with Tukey’s multiple-comparisons test). (I, J) Flow-cytometric analysis of tumor-infiltrating lymphocytes showing increased intratumoral CD8⁺ T-cell abundance following PSME4 depletion or PipAc2-4 pretreatment, whereas the measured regulatory T-cell population was not significantly altered. Data are presented as cell counts per mg tumor tissue (n = 10; one-way ANOVA with uncorrected Fisher’s LSD). (K) Survival of mice bearing control or PSME4-depleted tumors with or without anti-PD-1 treatment (Kaplan-Meier analysis; n = 9-10 per group; pairwise log-rank tests). (L) Survival of mice bearing control or PSME4-depleted tumors with or without IL-6 neutralization (Kaplan-Meier analysis; n = 7-8 per group; p values as indicated; pairwise log-rank tests).

To determine whether the restoration of tumor-intrinsic interferon competence extended beyond the *in vitro* setting, we evaluated STAT1 levels from the resected tumors. Immunoblot analysis demonstrated a fourfold increase in STAT1 abundance in PSME4-deficient tumors, whereas total STAT3 remained largely unchanged (**Fig. 6C** and **Fig. S8B**). Flow-cytometry analysis of syngeneic tumors showed higher CD8⁺ infiltration and favorable CD8-to-Treg/CD4 ratios in shPSME4 tumors (**Fig. S8C**). Differential proteomic analysis further revealed broad induction of interferon-associated proteins (74 of 85 significantly altered proteins in this pathway), accompanied by reduced abundance of proteins associated with IL-6 signaling, including STAT3 (9 of 11 significantly altered pathway components; **Fig. 6D**, **Fig. S8D,E** and **Supplementary Table S6**), supporting a shift toward an interferon-associated and away from an IL-6-associated tumor inflammatory state *in vivo*.

Because PipAc2-4 has not been optimized for systemic administration, we used *ex vivo* treatment to test whether transient PSME4 engagement, before tumor cell injection, could influence subsequent tumor-associated phenotypes. We performed a priming-rechallenge model (adapted from^41–44^). KP1.9 cells were injected into mice, fourteen days after priming, mice were rechallenged subcutaneously with KP1.9 shCtrl cells pre-treated with PipAc2-4 or DMSO, or with KP1.9 shPSME4 cells pre-treated with DMSO, and analyzed 48 hours thereafter (**Fig. 6E**). PipAc2-4 pretreatment and genetic PSME4 depletion each reduced tumor mass following rechallenge (**Fig. 6F**). Tumors arising under these conditions showed increased STAT1 abundance and reduced IL-6 secretion (**Fig. 6G-H**), indicating that transient PSME4 perturbation before implantation is sufficient to alter subsequent tumor phenotypes in this model. Both PSME4 depletion and PipAc2-4 pretreatment increased intratumoral CD8+ T-cell abundance, whereas the measured regulatory T-cell frequency was not significantly altered (**Fig. 6I-J** and **Fig. S8G**).

Tumor-intrinsic interferon signaling is a well-established prerequisite for effective immune checkpoint blockade^45–47^. We thus examined whether PSME4 status altered the response to PD-1 blockade. Under the treatment conditions used, anti-PD-1 did not significantly prolong survival in either control or PSME4-deficient tumors (**Fig. 6K** and **Fig S8H**). PSME4 depletion alone led to greater tumor reduction than the tested anti-PD-1 regimen, and addition of anti-PD-1 did not further improve the outcome. Thus, in this model, PSME4 depletion suppresses tumor progression without requiring efficacy from concurrent PD-1 blockade. Because the model showed limited sensitivity to anti-PD-1, checkpoint-responsive models will be required to determine whether PSME4 targeting can enhance or alter responsiveness to immune checkpoint therapy. Nevertheless, because PSME4 depletion consistently reduced IL-6 production and secretion across human and murine cell lines, we asked whether IL-6 neutralization could further affect tumor outcome. Previous studies have suggested that IL-6 signaling may represent a therapeutic target in NSCLC^48–50^, but its relationship to PSME4 has not been examined. Notably, anti-IL-6 treatment prolonged survival in mice bearing PSME4-deficient KP1.9 tumors but did not significantly benefit control tumors under the conditions tested (**Fig. 6L**), suggesting that PSME4 depletion creates a context in which residual IL-6 activity may be therapeutically exploited. Collectively, these findings show that PSME4 depletion stabilizes STAT1 *in vivo*, remodels the tumor immune microenvironment, increases CD8+ T-cell accumulation, and suppresses tumor growth. In the model examined here, these effects were not further enhanced by the tested anti-PD-1 regimen, whereas IL-6 neutralization preferentially benefited PSME4-deficient tumors. Together with our previous demonstration that PSME4 restricts antigen presentation, these results support complementary roles for proteasome composition in controlling tumor immunogenicity and tumor-intrinsic inflammatory competence.

## Discussion

By building a novel chemical biology toolkit, identifying and validating a selective covalent probe, and applying the probe in combination with genetics, our studies define a pathway regulated by the poorly understood proteasome activator PSME4 whose immune functions were previously attributed to MHC-I antigen presentation. Unexpectedly, we found that PSME4 specifically regulates the STAT1-IL-6/STAT3 pathway. Upregulation of PSME4 in NSCLC makes this an attractive candidate for inhibition, and indeed pharmacological inhibition of this process with our probe suggests a route to preventing tumor growth.

Although tumor-intrinsic interferon signaling has long been known to be regulated by the ubiquitin system, our finding a role of PSME4 reveals a key role for ubiquitin-independent proteasomal degradation. Loss of interferon responsiveness in cancer has been linked to genetic alterations in IFN receptors, JAK kinases, and downstream signaling components, as well as defects in upstream innate immune pathways^38,51–53^. Our findings suggest that altered turnover of a central signaling effector can attenuate the same transcriptional program without requiring genetic disruption of the pathway. PSME4-dependent STAT1 turnover may therefore provide an endogenous, non-genetic mechanism for tuning the amplitude of tumor-intrinsic interferon signaling. This is particularly relevant because both impaired IFN responsiveness and sustained IFN signaling have been associated with resistance to immune checkpoint blockade, suggesting that the magnitude and duration of this pathway must be tightly controlled. This distinction may also provide a mechanistic connection between proteasome composition and the reciprocal behavior of interferon-associated and IL-6/STAT3-associated programs observed across tumors^54,55^. Understanding of how STAT1 stabilization is coupled with reduced STAT3 activity and IL-6 production remains a question for the future. Nonetheless, changes in STAT-containing complexes following PSME4 depletion suggest one potential point of interaction, but transcriptional feedback and additional PSME4-regulated substrates are also likely to contribute. Likewise, although our degradome, proteasome-association, and pharmacological data support PSME4-dependent STAT1 turnover, the molecular basis of substrate recognition remains unknown. While PSME4 is thought to act in a ubiquitin-independent manner^16,17,56^, biochemical reconstitution will be required to determine whether STAT1 is directly recognized and degraded by PSME4-capped proteasomes. Whether recognition depends on a particular post-translational or conformational state or specific subcellular localization, and whether additional ubiquitin-dependent mechanisms contribute to STAT1 turnover, remains to be elucidated.

An important aspect of this study is the ability to pharmacologically perturb a non-catalytic proteasome regulator. Conventional proteasome inhibitors target catalytic sites within the 20S core and broadly suppress proteasomal degradation. By contrast, covalent engagement of PSME4 C1001 altered selected proteasome activities and reproduced established consequences of PSME4 depletion without globally inhibiting the proteasome core. Our structural analysis provides a mechanistic basis for this selectivity. Previous structures established the overall architecture of the PSME4-20S complex, including PSME4-dependent remodeling of the 20S gate and the presence of apertures proposed to provide routes for substrate entry or product release^8,32,33^ . The PipAc2-4-bound structure remains largely consistent with this overall architecture but reveals a distinct local configuration around C1001, adjacent to a region proposed to mediate substrate entry or product release. Although the functional consequences of these structural changes remain to be established, their location raises the possibility that covalent engagement at C1001 modulates substrate access, processing, or product transit rather than simply suppressing 20S catalytic activity. This model is consistent with the selective changes in proteasome activity observed following PipAc2-4 treatment and provides a structural framework for understanding how liganding a non-catalytic regulator can reshape proteasome function without globally inhibiting the core particle.

The chemical proteomic strategy used to discover PSME4 ligands has broader implications. Reducing electrophile reactivity through ring expansion did not simply decrease proteome-wide engagement but uncovered selective ligand-protein interactions that were weak or absent with the more reactive azetidine acrylamide scaffold, suggesting that tempering intrinsic reactivity can favor the identification of covalent probes that depend more on molecular recognition, including those that target previously undrugged proteins. Here, we reveal how matching a probe to a target can fuel biological perspective. From a biological perspective, the impact of PipAc2-4, remodeling of proteasome activity by chemically targeting a regulatory component, provides a distinct strategy from both catalytic proteasome inhibition, which leads to a general blockade of protein turnover, and targeted protein degradation approaches that promote destruction of selected proteins. Indeed, we show that PSME4 ligands invert the conventional targeted-degradation paradigm in that pharmacological manipulation of the degradation machinery is used to preserve rather than eliminate a regulatory protein. Whether this principle can be extended to other proteasome regulators or multiprotein degradation complexes remain to be determined.

With a specific pathway controlled by PSME4 in NSCLC models elucidated, we note that PSME4 expression is altered across cancer contexts. Whether PSME4-dependent STAT1 turnover represents a general mechanism or is determined by tissue state, genotype, or proteasome composition remains an important question for the future. Functions of PSME4 in normal tissues will also require reconsideration as PSME4-directed pharmacology advances. More broadly, the additional candidate substrates identified by degradome profiling suggest that the consequences of PSME4 activity are unlikely to be restricted to STAT1 and may vary across cellular contexts. Defining these substrate repertoires will be important for understanding how individual proteasome assemblies acquire functional specificity.

Our findings, therefore, support a broader view of proteasome composition in which expression of specific regulatory components influences not only the products of proteolysis but also the selection of proteins committed to degradation. In tumors, PSME4 illustrates how this layer of proteostasis can simultaneously influence antigen presentation and inflammatory signaling, connecting protein degradation to both immune recognition and tumor cell state. Together, our current and previous findings position proteasome composition as a regulatory layer that controls both what tumor cells present to the immune system and how they interpret inflammatory signals. The ability to selectively perturb a non-catalytic proteasome regulator further establishes proteasome composition as an experimentally and pharmacologically accessible dimension of proteostasis, with implications extending beyond tumor immunity.

## Limitations

PipAc2-4 represents a chemical probe rather than an optimized therapeutic candidate. Its current pharmacological properties preclude systemic administration, restricting its use to mechanistic cellular studies and *ex vivo* perturbation. Further optimization of potency, solubility, pharmacokinetics, and *in vivo* target engagement will therefore be required to determine the therapeutic potential of pharmacological inhibition of PSME4. The *in vivo* studies also highlight both the potential and the current boundaries of this strategy. Genetic PSME4 depletion suppressed tumor growth and altered the immune microenvironment, but the KP1.9 model showed limited sensitivity to PD-1 blockade and therefore does not establish whether PSME4 targeting can enhance checkpoint responsiveness. Similarly, the preferential benefit of IL-6 neutralization in PSME4-deficient tumors identifies a context-dependent vulnerability but does not yet establish PSME4 as a predictive biomarker for IL-6-directed therapy. Given previous efforts to therapeutically target IL-6 signaling in cancer, including NSCLC, our findings raise the possibility that tumor inflammatory state or PSME4 status could help define contexts in which IL-6-directed intervention is more effective, although this will require testing across additional models. Testing pharmacologically optimized PSME4 inhibitors in checkpoint-responsive models, as well as in combination with interventions that directly potentiate interferon signaling, will be important for defining how these inflammatory states can be therapeutically exploited.

## Supporting information

Data Set S1

## Extended Data Figure legends

**Fig. S1** | **Glutathione (GSH) reactivity and cysteine-directed ABPP of ring-expanded stereoprobes.** (A) GSH reactivity of ring-expanded stereoprobes. AzAc reactivity was assessed at 2 h, PyAc reactivity at 2 and 6 h, and PipAc reactivity at 6 h. Each stereoprobe is represented as an individual point and grouped according to ring size. Data represents mean values from 2-3 independent experiments. (B) Volcano plots of cysteine-directed ABPP data from 22Rv1 cells treated with cis-AzAc, PyAc, PipAc1, or PipAc2 stereoprobes (20 μM, 3 h). Cysteines showing ≥66.7% reduction in IA-DTB labeling relative to DMSO and ≥2.5-fold stereoselectivity between enantiomeric pairs are highlighted in red or blue (two-tailed Student’s t test). Data represents average values from 4-6 independent experiments.

**Fig. S2 | Stereoselective engagement of recombinant PSME4 by PipAc probes.** (A) Left, gel-ABPP showing stereoselective engagement of recombinant wild-type PSME4 expressed in HEK293T cells by the indicated PipAc1-yne stereoprobes. Right, full-range gel image corresponding to the same experiment. Cells were treated with PipAc1-yne stereoprobes (10 μM, 3 h), lysed, and stereoprobe-modified proteins were conjugated by copper-catalyzed azide-alkyne cycloaddition (CuAAC) to a rhodamine-azide reporter, separated by SDS-PAGE, and visualized by in-gel fluorescence. Mock-transfected cells served as vector controls. Data shown are from one experiment representative of three independent experiments. (B) Sequence conservation of PSME4 across vertebrate species. PSME4 orthologs from *Danio rerio* (zebrafish), *Xenopus laevis* (frog), *Mus musculus* (mouse), *Rattus norvegicus* (rat), *Macaca fascicularis* (cynomolgus macaque), and *Homo sapiens* (human) were aligned using Clustal Omega. The displayed region corresponds to residues 975-1025 of human PSME4. Conserved residues are shown in yellow, with C1001 highlighted; the consensus sequence is shown above the alignment. (C) Cysteine-directed ABPP showing stereoselective engagement of the PTGES2 C110/C113-containing peptide by PipAc1-2 (left), but not PipAc2-4 (right), in 22Rv1 cells. Data represent mean ± SEM from 4-6 independent experiments. (J-L) Comparison highlighting side chain rearrangements of the PipAc2-4-bound pocket of PSME4 with previously determined structures: (J) 6REY (salmon), (K) 6KWY (yellow), and (L) 7NAQ (blue). (M) Superimposition of PipAc2-4-bound pocket with all three previously determined models.

**Fig. S3 | Cryo-EM Processing Scheme for PipAc2-4-bound PSME4-20S CP complex.(A)** Representative micrograph (scale bar: 300Å) and selected 2D classes (scale bar: 170Å for subsequent refinements. **(B)** 3D reconstruction of the PipAc2-4-bound PSME4-20S CP complex at 2.38 Å, along with local resolution projected on a sharpened volume. **(C)** Gold-standard Fourier shell correlation (GSFSC) plot (top panel), and angular distribution plot (bottom panel) of particles from final refinement of the PipAc2-4-bound PSME4-20S CP complex. Mask used for local refinement is shown in cyan. **(D)** Focused refined 3D reconstruction on PSME4 along with local resolution projected on a sharpened volume. **(E)** Gold-standard Fourier shell correlation (GSFSC) plot (left panel) and angular distribution plot (right panel) of particles from local refinement of the PipAc2-4-bound PSME4-20S CP complex.

**Fig. S4 | Comparison of PipAc2-4-PSME4-20S CP with previously determined Cryo-EM structures and maps.** (A-C) Comparison highlighting side chain rearrangements of the PipAc2-4-bound pocket of PSME4 with previously determined structures: (A) 6REY (salmon), (B) 6KWY (yellow), and (C) 7NAQ (blue). Superimposition of PipAc2-4-bound pocket with all three previously determined models presented in Fig. 2G. (D) Structural details of the putative opening as observed in PipAc2-4-bound PSME4-20S CP. The loops surrounding the opening are colored in yellow (loop A), cyan (loop B), and purple (loop C). Comparison highlighting side chain rearrangements of the PipAc2-4-bound pocket of PSME4 with previously determined structures: 6REY (salmon), 6KWY (yellow), and 7NAQ (blue). (E) Comparison of Cryo-EM maps at opening-1 of PipAc2-4-bound PSME4 (green), 6REY (salmon), 6KWY (yellow), 7NAQ (blue).

**Fig. S5 | Biochemical selectivity of PipAc2-4 for** β**1-associated proteasome activity and validation of reagents.** (A, B) Purification and verification of 20S-PSME4 complex. Coomassie-stained gel and immunoblots confirm the composition and integrity of purified 20S-PSME4 complexes, with PSME4 detected in the E1/E2 fractions. (C-E) Proteasome activity assay in time courses with the stereoisomer panel: PipAc2-3 (C), PipAc2-1 (D), and PipAc2-2 (E) at 1, 10, 100 μM. PipAc2-4 produces a clear, dose-dependent reduction in nLPnLD-AMC cleavage; other stereoisomers show minimal effects. (F-G) Dose-response traces for fluorogenic proteasome substrates across increasing ligand concentrations 1, 10, 100 μM (log₂ scale): (F) β5 (LLVY-AMC; chymotrypsin-like), and (G) β2 (RLR-AMC; trypsin-like) (n=2). (H-I) Endpoint analyses (3 h) of caspase-like activity in 20S versus 20S-PSME4 proteasomes (nLPnLD-AMC substrate) presented in percent out of 20S-PSME4 activity; MG132 serves as a control. (I) 20S-PSME4 proteasomes with PipAc2-4 treatment or DMSO as control, showing reduced β1-associated nLPnLD-AMC cleavage presented in percent out of 20S-PSME4 DMSO-treated activity. (J-K) Endpoint analyses (3 h) of A549-derived proteasomes following 18 h treatment with PipAc2-4 or DMSO show reduced β1-associated nLPnLD-AMC cleavage without significant effects on β5 (LLVY-AMC) or β2 (RLR-AMC) activities (mean ± SEM n = 4; p values as indicated). (L-N) Validation of cellular perturbations. (L) Immunoblots showing expression of Luc (Luciferase), PSME4(WT), or PSME4(C1001A) in A549 PSME4-KO cells; blotted for PSME4, His-tag, and Actin (loading control). (M,N) Immunoblots confirming efficient PSME4 knockdown in A549 (M) and KP1.9 (N) cells using shPSME4 relative to shCtrl.

**Fig. S6 | Additional degradome and biochemical analyses supporting PSME4-dependent STAT1 regulation.** (A, B) Heat maps of calculated Z scores for peptides (A) or proteins (B) differentially represented in MAPP analysis of A549 cells treated with PipAc2-4 (10 μM, 18 h) or DMSO, showing relative abundance across replicates. (C, D) Heat maps of calculated Z scores for peptides (C) or proteins (D) differentially represented following PSME4 overexpression relative to GFP control. (E) Venn diagram showing overlap among peptides translated into proteins with altered degradation following PipAc2-4 (DMSO) and PSME4 overexpression, identifying 60 shared proteins (16.9%), including STAT1. (F) Immunoblot analysis of STAT1 and STAT3 abundance in control and PSME4-depleted A549 cells following bortezomib treatment.

**Fig. S7 | Additional analyses of PSME4-dependent cytokine regulation.** (A) Independent STAT3 immunoprecipitation under non-denaturing conditions showing increased STAT1 co-association following PSME4 depletion. (B) Independent immunoblot analysis of A549 cells showing reduced STAT3 phosphorylation relative to total STAT3 following PSME4 depletion, with total STAT3 abundance largely unchanged. (C) Multiplex cytokine profiling by Luminex in KP1.9 tumors expressing shCtrl or shPSME4. Cytokine abundance is presented as the shPSME4/shCtrl ratio across all measured analytes. (D) Multiplex cytokine profiling following 6 h. TNFα/IFNγ stimulation in control and PSME4-depleted cells, highlighting selected cytokines, including reduced IL-6. (E) IL-6 ELISA following treatment with the inactive PipAc2 stereoisomers PipAc2-1, PipAc2-2, or PipAc2-3 (1-50 μM), showing no comparable suppression of IL-6 secretion relative to the active PipAc2-4 probe.

**Fig. S8 | Additional in vivo analyses of PSME4-dependent tumor inflammation and immune composition.** (A) Cell proliferation analysis of KP1.9 shCtrl and shPSME4 cells using a Cell Counting Kit-8 assay, showing no significant difference in *in vitro* proliferation between the two lines. (B) Quantification of the immunoblot shown in Fig. 6C for PSME4, STAT1, and STAT3 abundance in control and PSME4-depleted tumors, normalized to actin (mean ± SEM, n = 3; Student’s t test). (C) Flow-cytometric analysis of syngeneic tumors showing CD8⁺ T-cell abundance normalized to tumor mass and the indicated CD8 and CD8 ratios following PSME4 depletion. (D) Differential proteomic analysis of control and PSME4-depleted tumors, highlighting PSME4/STAT1-associated changes *in vivo*. (E) Bubble plot showing enriched pathways identified inshPSME4 tumors using STRING Local Network Clusters. Bubble size indicates the number of proteins contributing to each pathway, and color denotes the Benjamini-Hochberg-adjusted p value (FDR). (F) Flow-cytometric analysis of the immune composition of priming-challenged syngeneic tumors, expressed as a percentage of CD45⁺ cells, showing changes in macrophage and NK1.1⁺ populations following PSME4 depletion, while total CD45⁺ and CD11b⁺ populations remain comparable, as indicated. (G) Representative flow-cytometry gating strategy used to identify tumor-infiltrating immune populations quantified in the study. (H) Representative H&E-stained lung sections from mice injected with KP1.9 control or PSME4-depleted cells and treated with control IgG or anti-PD-1 antibody. A representative lung from a non-injected mouse is shown for comparison. Scale bars, 2,000 μm.

## Abbreviations and Symbols

20S: 20S core proteasome.
26S: 26S proteasome.
ABPP: activity-based protein profiling.
AzAc: azetidine acrylamide stereoprobe.
B2M: β2-microglobulin.
CuAAC: copper(I)-catalyzed azide-alkyne cycloaddition.
DMSO: dimethyl sulfoxide.
FDR: false discovery rate.
GFP: green fluorescent protein.
GSH: glutathione.
HLA-A/B/C: human leukocyte antigen class I molecules.
IA-DTB: iodoacetamide-desthiobiotin.
IFI44: interferon-induced protein 44.
IFIT1 / IFIT2: interferon-induced proteins with tetratricopeptide repeats 1 and 2.
IFN: interferon.
IL-6: interleukin-6.
IRF1: interferon regulatory factor 1.
ISG: interferon-stimulated gene.
ISG15: interferon-stimulated gene 15.
JAK-STAT: Janus kinase-signal transducer and activator of transcription.
JNK: c-Jun N-terminal kinase.
KO: knockout.
MAPP: mass spectrometry-assisted proteasome profiling.
MFI: mean fluorescence intensity.
MHC-I: major histocompatibility complex class I.
MS: mass spectrometry.
NF-κB: nuclear factor kappa B.
NSCLC: non-small-cell lung cancer.
OE: overexpression.
PD-1: programmed cell death protein 1.
PipAc: piperidine acrylamide stereoprobe.
PSMA1: proteasome 20S subunit alpha 1.
PSMB2: proteasome 20S subunit beta 2.
PSME4 / PA200: proteasome activator complex subunit 4 / proteasome activator 200 kDa.
PTGES2: prostaglandin E synthase 2.
PyAc: pyrrolidine acrylamide stereoprobe.
qPCR: quantitative polymerase chain reaction.
SDS-PAGE: sodium dodecyl sulfate-polyacrylamide gel electrophoresis.
SEM: standard error of the mean.
shCtrl: control short hairpin RNA.
shPSME4: short hairpin RNA targeting PSME4.
STAT1 / STAT3: signal transducer and activator of transcription 1 and 3.
TNFα: tumor necrosis factor alpha.
TNFAIP3: tumor necrosis factor alpha-induced protein 3.
Treg: regulatory T cell.
WT: wild-type.
β1 / β2 / β5: proteasome catalytic subunits.
β1i / LMP2: immunoproteasome catalytic subunit β1i

## Acknowledgments

We thank the members of the Merbl, Cravatt, and Schulman laboratories for their support. We thank Daniel Bollschweiler and Tillman Schäfer of the Max Planck Institute of Biochemistry Cryo-EM facility (RRID:SCR_025744) for their assistance with data collection as well as J. Rajan Prabu for maintaining processing infrastructure. We thank Sussane V Gronau for baculoviral Insect cell expressions. We thank Jakob Farnung and Samuel Maiwald for fruitful and scholarly discussions. This work was supported by the Israel Science Foundation (grant no. 2237/23) and by the European Union: ERC CoG (grant no. 101045613) and ERC PoC (grant no. 101335588). Views and opinions expressed are, however, those of the author(s) only and do not necessarily reflect those of the European Union or the European Research Council. Neither the European Union nor the granting authority can be held responsible for them. The Merbl lab is supported by the Cancer Grand Challenges partnership funded by Cancer Research UK (ILLUMINE Team, CGCATF-2025/100036); the Lotte and John Hecht Memorial Foundation; the Knell Family Institute for Artificial Intelligence. Y.M. is a CRI Lloyd J. Old STAR (CRl5602), a CRI/Israel Cancer Research Fund Technology Impact Award recipient (CRI5398) and a Cancer Research Institute/Israel Cancer Research Fund CLIP Grant (CRI4351). A.L. is supported by the Fellowship of the Center for Integration in Science, Israel Ministry of Aliyah and Integration, K.G. and R.S. are supported by Clinical Co-Mentoring fellowships through the Moross Integrated Cancer Center. The Schulman lab is supported by Max Planck Society.

