## Supplemental Figures for "Targeting proteasome composition reprograms tumor-intrinsic inflammation through selective degradation of STAT1"

**Fig. S1 | Glutathione (GSH) and proteomic reactivity of ring-expanded stereoprobes.**

**A**

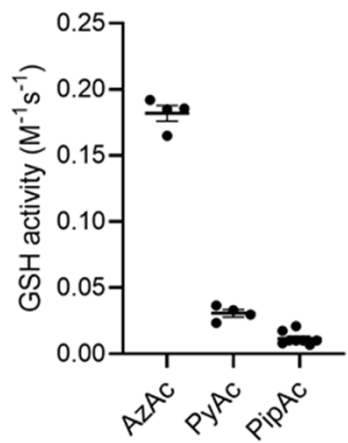

**B**

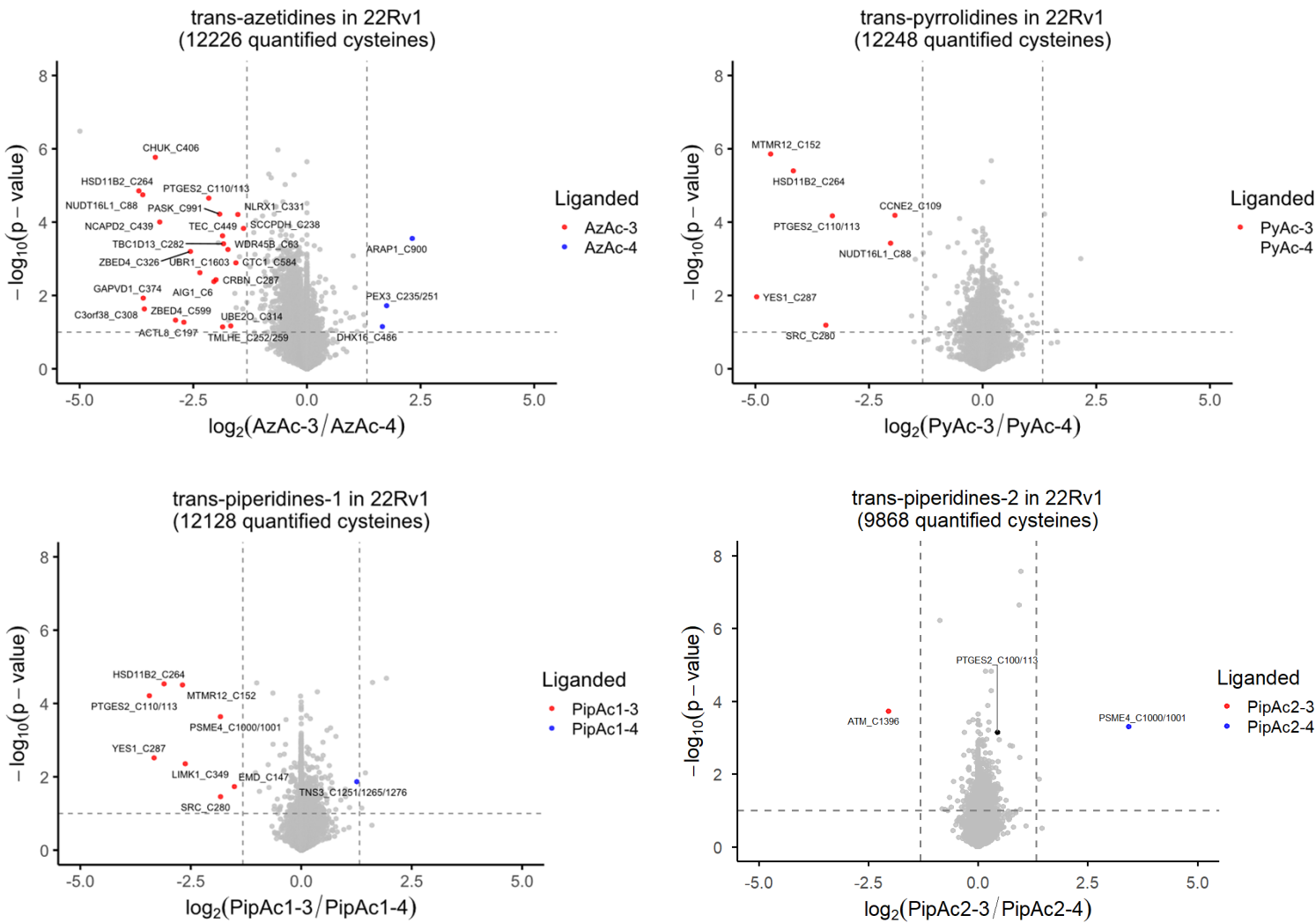

Fig. S2 | Stereoselective engagement of recombinant PSME4 by piperidine-1 alkyne probes.

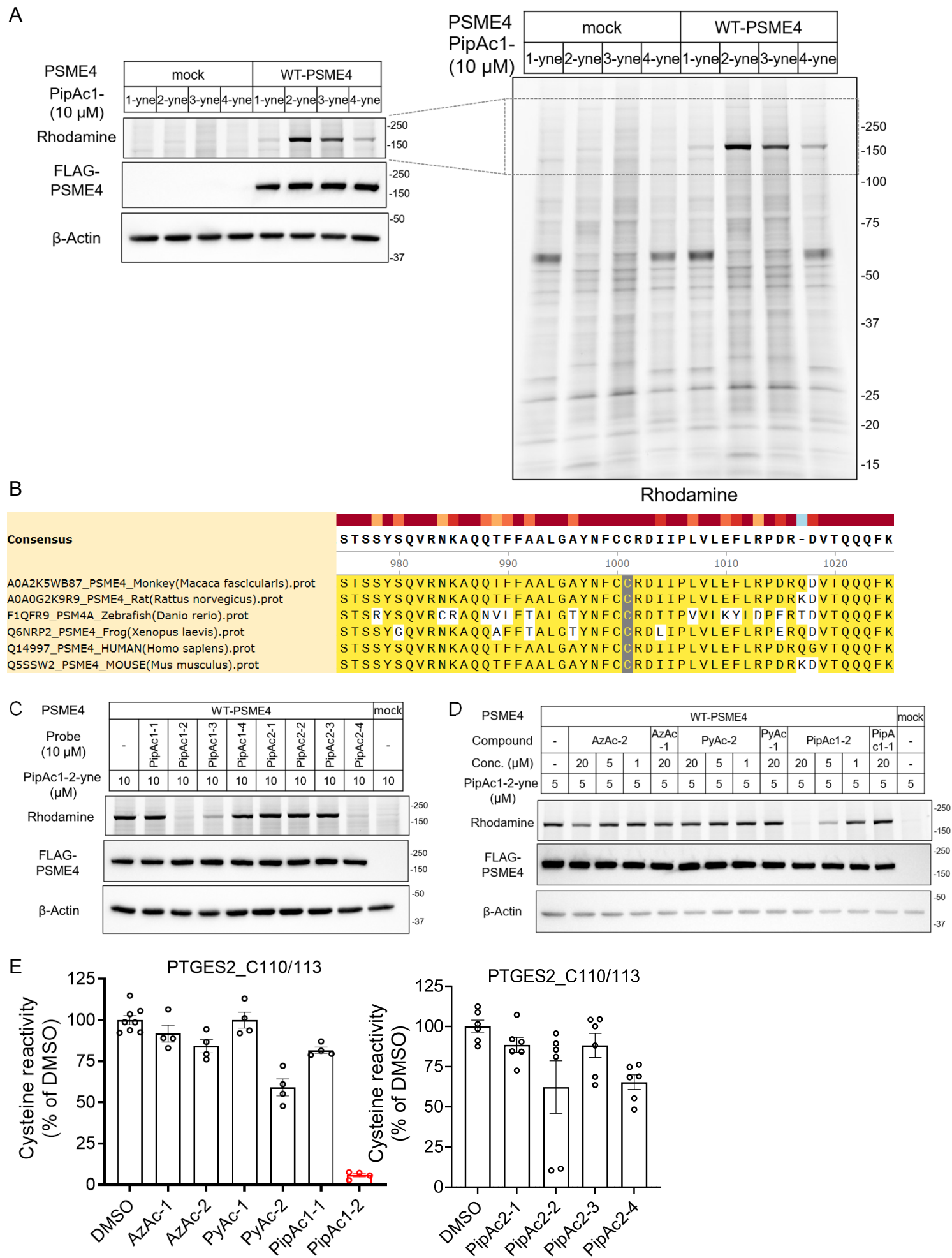

**Fig. S3 | Cryo-EM Processing Scheme for PipAc2-4-bound PSME4–20S CP complex.**

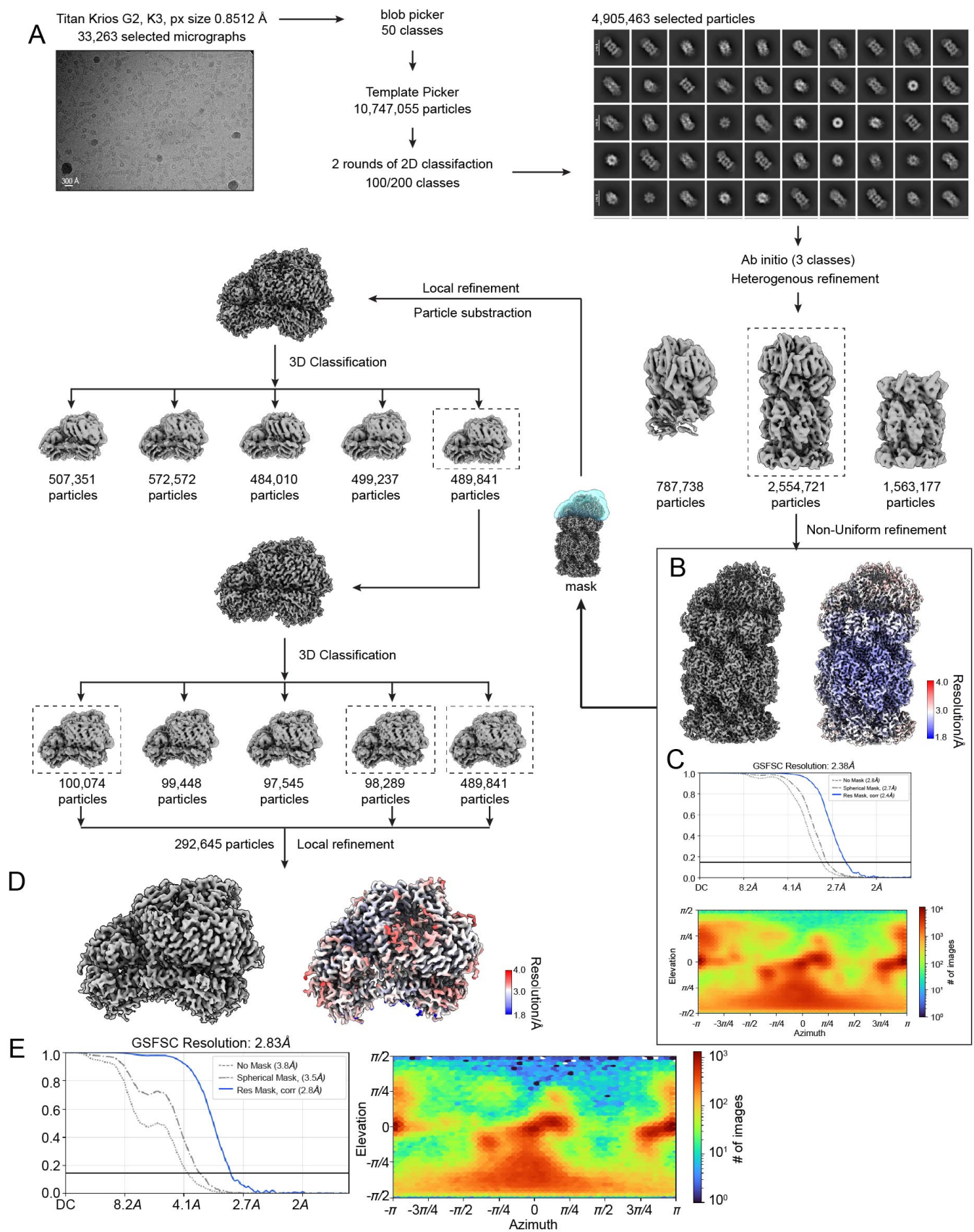

**Fig. S4 | Comparison of PipAc2-4-PSME4-20S CP with previously determined Cryo-EM structures and maps.**

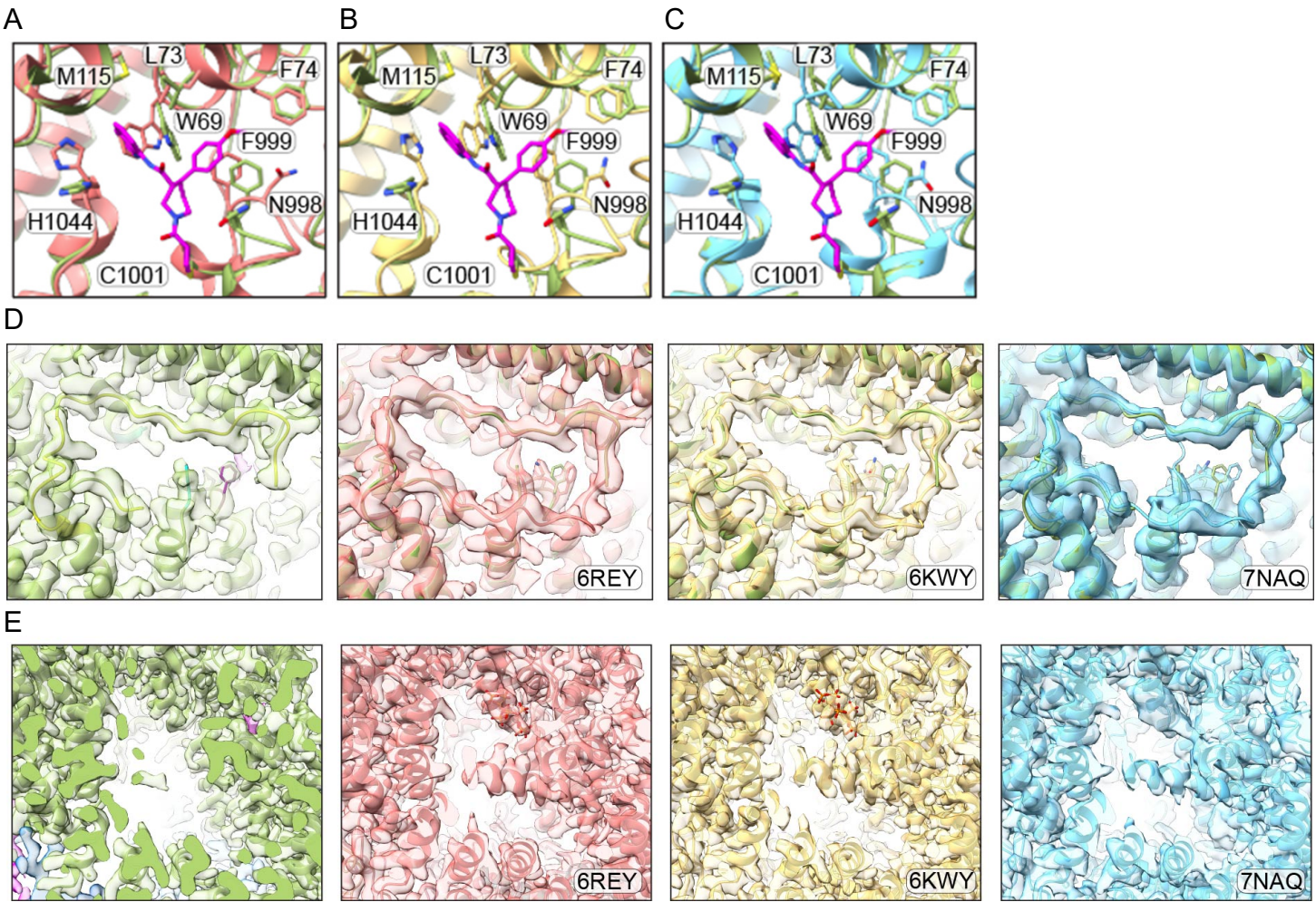

**Fig. S5 | Biochemical selectivity of PipAc2-4 for β1-associated proteasome activity and validation of reagents**

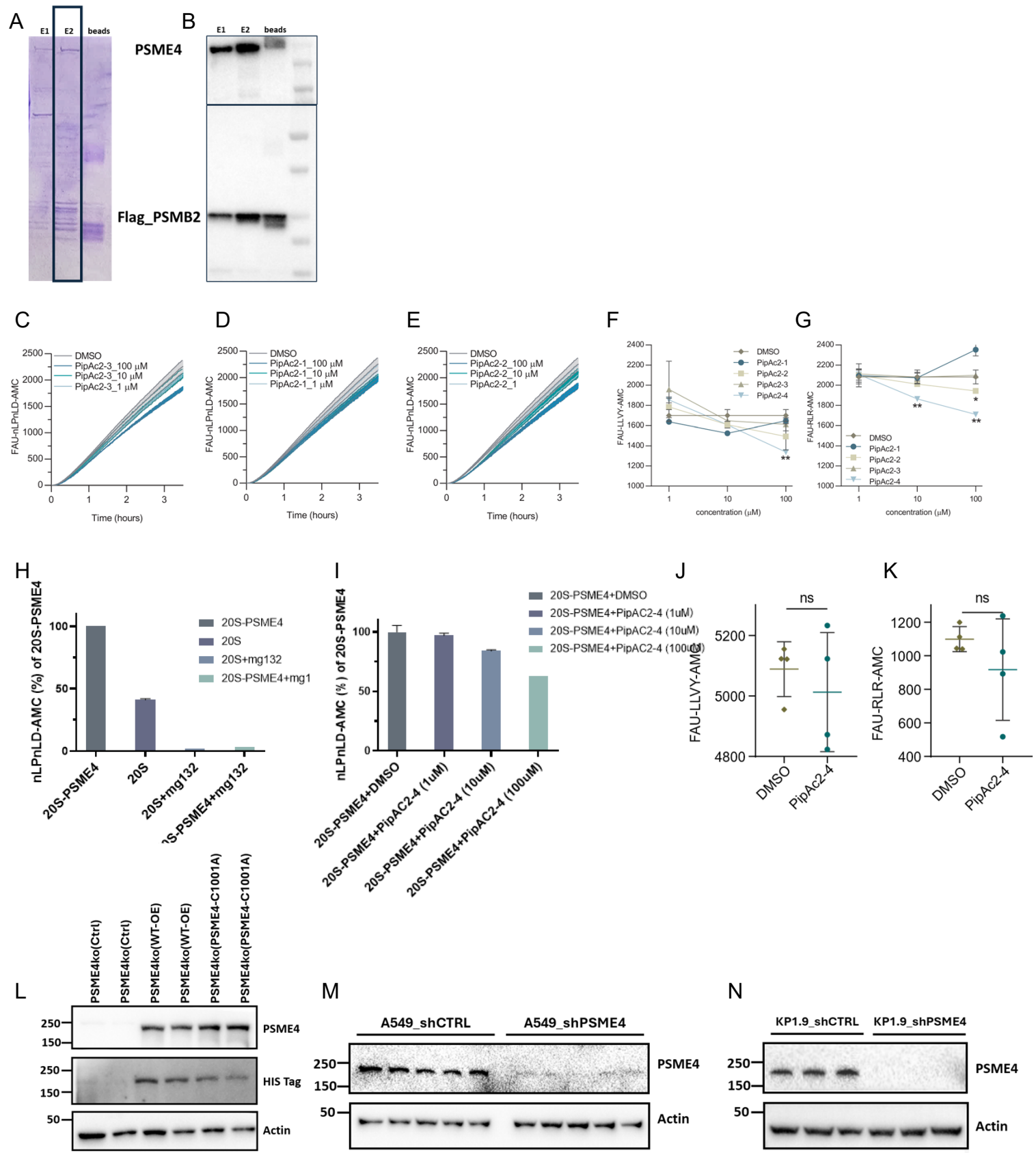

**Fig. S6 | Additional degradome and biochemical analyses supporting PSME4-dependent STAT1 regulation.**

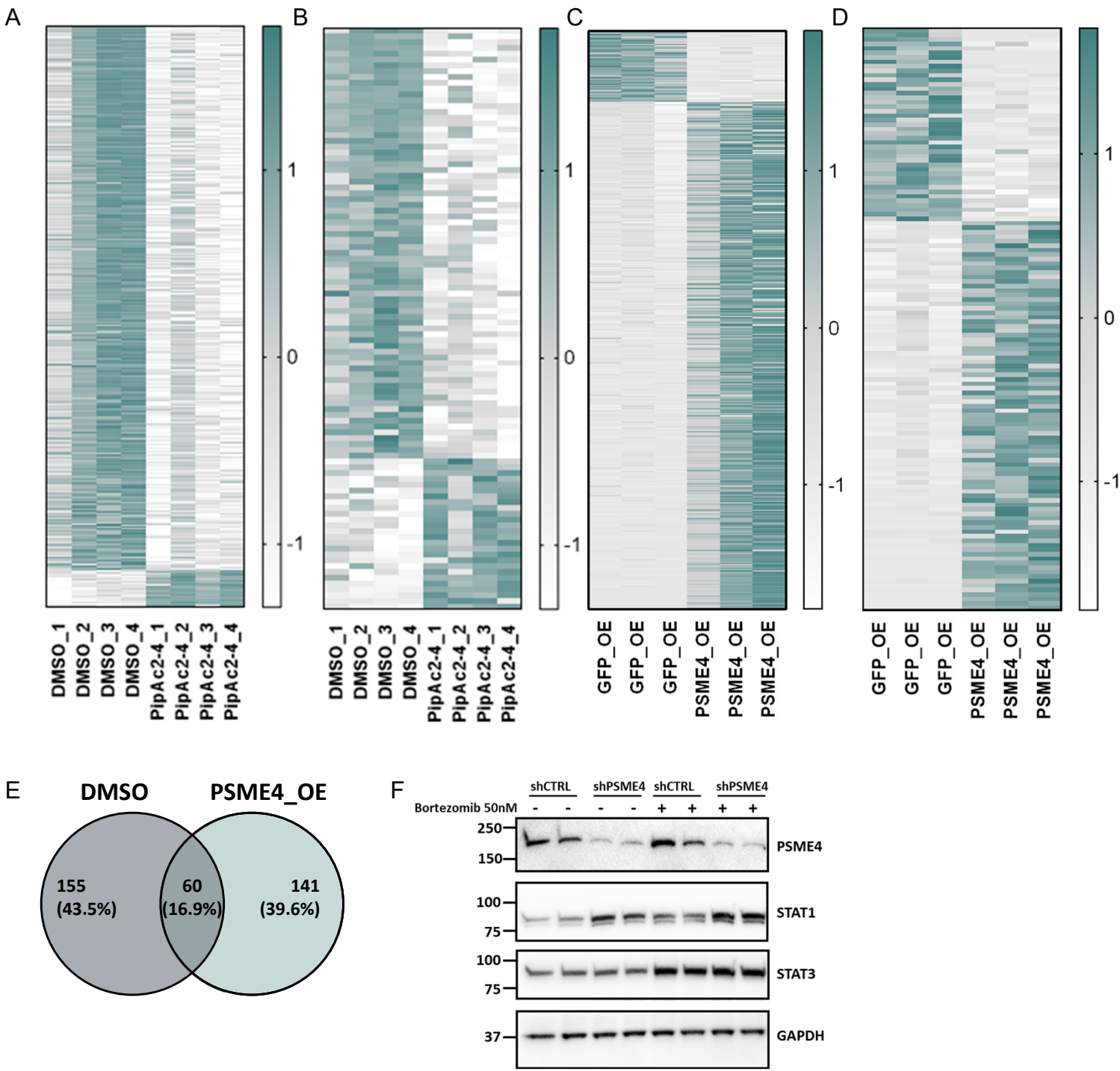

Fig. S7 | Additional analyses of PSME4-dependent STATs and cytokine regulation

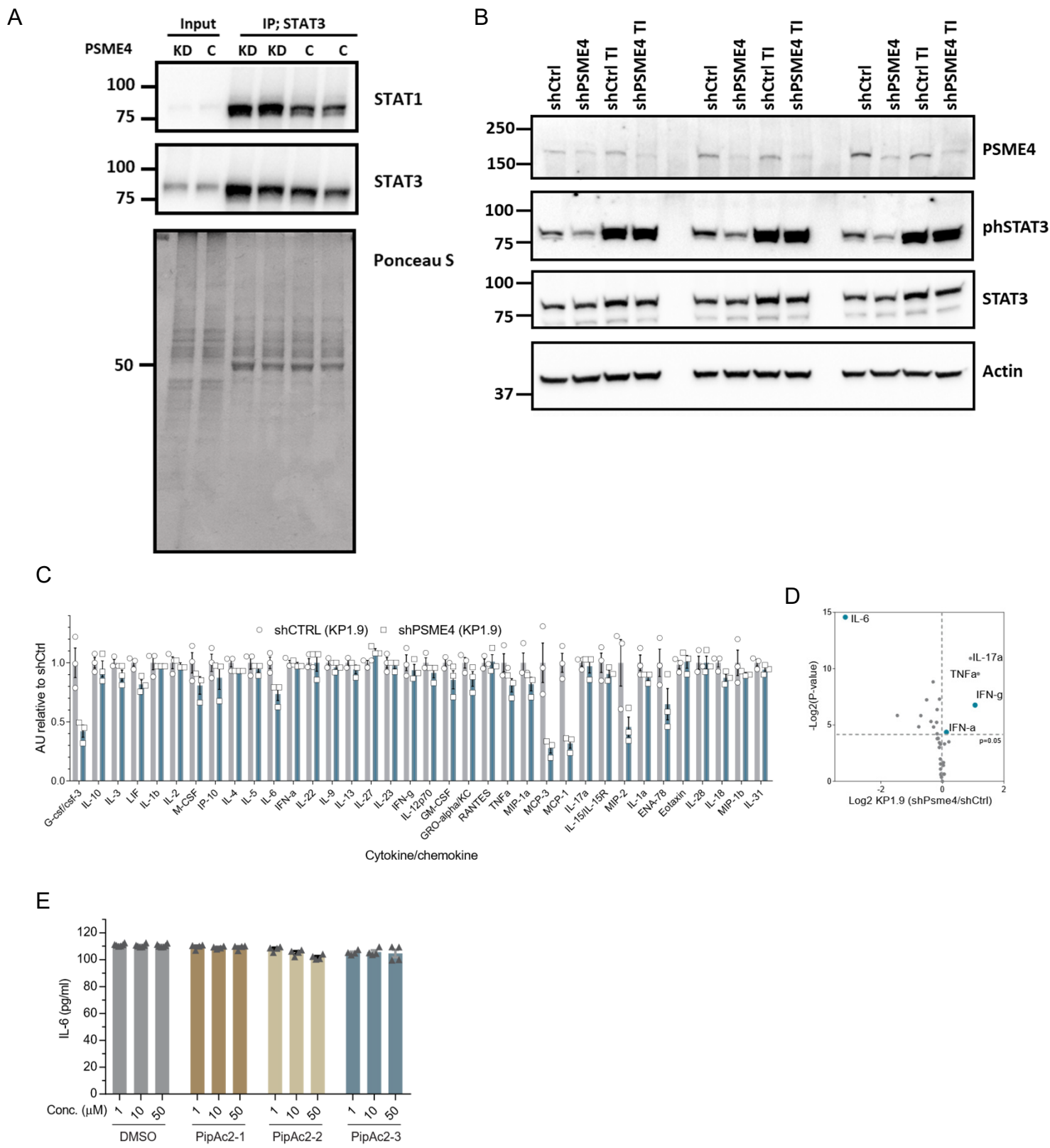

**Fig. S8 | Additional in vivo analyses of PSME4-dependent tumor inflammation and immune composition.**

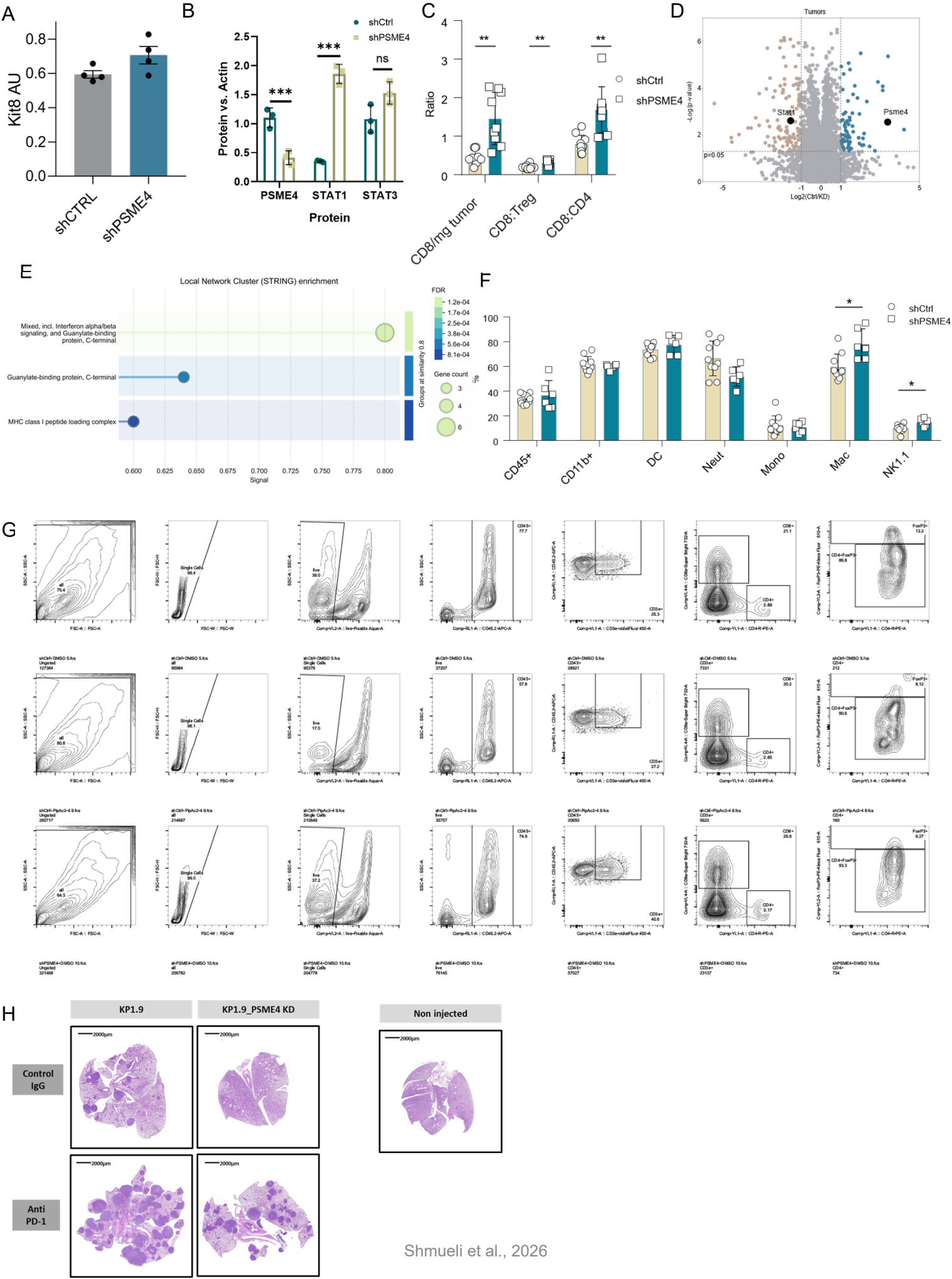
