## Supplementary material for "Targeting proteasome composition reprograms tumor-intrinsic inflammation through selective degradation of STAT1": Materials and Methods

**Gel-based activity based-protein profiling**

HEK293T cells (10 cm dish, 60% confluency) were transiently transfected with FLAG-PSME4 or empty vector (mock) for 24 h then seeded to 6-well plate (3 X105 cells/well) and grown for 16 h to ~50% confluency. Cells were then treated with DMSO or stereoprobes in 1.5 mL complete media for the indicated time and concentration, followed by alkyne probes in 1.5 mL complete media for 1 h and the indicated concentration before harvesting. Pellets were lysed in DPBS by sonication (40%, 2 X 12 pulses), clarified by centrifugation (16,000g for 10 min at 4 °C), and adjusted to 1 mg/mL total protein concentration. Copper-catalyzed azide-alkyne cycloaddition (CuAAC) was performed by adding click-master mix (1/10 volume containing 1 μL 1.25 mM TAMRA-azide, 1 μL 50 mM CuSO4, 3 μL 1.7 mM TBTA in 1:4 DMSO: t-butanol, 1 μL 50 mM TCEP) and incubated for 1 hour at room temperature. The samples were then quenched with 4X Laemmli sample buffer, heated at 95 °C for 5 min, resolved by 4-20% Novex Tris-Glycine mini gels (Invitrogen) and scanned for fluorescent signals using a Bio-Rad Chemidoc imager. The band intensities were quantified using ImageJ and not normalized to loading controls.

**Cysteine-directed activity based-protein profiling**

Cysteine-directed ABPP was performed as previously reported^1^. In brief, 22Rv1 cells (20 million per 15 cm dish) or Ramos cells (12.5 mL at 2 million cells/mL) were treated with DMSO or stereoprobes (20 μM, 3 h). Cells were washed 3 times with ice-cold DPBS, resuspended in DPBS, and lysed by probe sonication (2 × 15 pulses; 10% output). The proteome was adjusted to 2 mg/mL in 500 μL and treated with 0.1 mM iodoacetamide-desthiobiotin (IA-DTB) for 1 hour at room temperature. Proteins were precipitated by adding ice-cold solvents (600 μL methanol, 200 μL chloroform, 100 μL water). After vigorous vortexing and centrifugation (16,000 × g, 10 min), solvents were removed by aspiration. The resulting protein disk was washed with 1 mL MeOH and resuspended in 90 μL reducing buffer (9 M urea, 10 mM DTT, 50 mM triethylammonium bicarbonate (TEAB), pH 8.5). Samples were heated (65 °C, 20 min), then alkylated with 50 mM iodoacetamide (30 min, 37 °C, in the dark). Samples were diluted to 2 M urea using 50 mM TEAB. Mass-spectrometry-grade trypsin (Promega; 1 μg in resuspension buffer with 25 mM CaCl2) was added and digestion proceeded overnight at 37 °C. Samples were incubated with streptavidin-agarose beads (50 μL slurry in 300 μL wash buffer containing 50 mM TEAB, 150 mM NaCl, 0.2% NP-40) for 2 hours at room temperature. Beads were transferred to BioSpin (Bio-Rad) columns, washed sequentially (3 × 1 mL wash buffer, 3 × 1 mL DPBS, 3 × 1 mL water), and enriched peptides were eluted with 300 μL elution buffer (50% acetonitrile, 0.1% formic acid) and dried by SpeedVac. Peptides were reconstituted in 200 mM EPPS (pH 8.0) and labeled with Tandem Mass Tag (TMT; 10-plex or 16-plex) for 90 min at room temperature. Labeled peptides were desalted using Sep-Pak C18 cartridges (Waters), fractionated by high-pH HPLC into 12 fractions, and analyzed on an Orbitrap Fusion mass spectrometer with Xcalibur v4.3.

**TMT liquid chromatography-mass-spectrometry (LC-MS) analysis**

All samples in this study used the same instruments and workflow settings as previously reported^1,2^. Briefly, samples were analyzed by Orbitrap Fusion Tribrid Mass Spectrometer (Thermo Scientific) coupled to an UltiMate 3000 Series Rapid Separation LC system and autosampler (Thermo Scientific Dionex). Data was acquired using an MS3-based TMT method^2^ (see ref for detailed settings) and the RAW files were uploaded and processed by Integrated Proteomics Pipeline (IP2, v.6.7.1), and searched using the ProLuCID algorithm (10.1016/j.jprot.2015.07.001) using a reverse concatenated, non-redundant variant of the Human UniProt database (release 2016). Cysteine residues were searched with a static modification for carboxyamidomethylation (+57.02146 Da) and a dynamic modification for IA-DTB labeling (+398.25292 Da) was included for a maximum number of 2 differential modifications. N-term and lysine static modification from the tags (+229.1629 Da for 10-plex, +304.2071 Da for 16-plex) were included. Peptides were required to be at least 6 amino acids long and filtered to through DTASelect (version 2.0) to obtain a false-positive peptide rate below 1%. The MS3-based peptide quantification was performed with reporter ion mass tolerance set to 20 ppm with Integrated Proteomics Pipeline (IP2).

**Data processing of cysteine-directed ABPP**

The census output files from Integrated Proteomics Pipeline 2 (IP2, v.6.7.1) were further processed to calculate cysteine engagement ratios (probe vs DMSO) by dividing each TMT reporter ion intensity by the average intensity for the DMSO channels. Peptide-spectra matches were grouped based on protein ID and the cysteine residue number. Peptides with summed reporter ion intensities < 10,000 or coefficient of variation for DMSO channels > 0.5 were excluded from analysis. TMT ion intensities were median normalized per TMT channel. Two or three independent replicates were analyzed together, and data were averaged from all replicates. Quantified cysteines located on the same tryptic peptide were combined. A cysteine site was considered stereoselectively liganded if the variability corresponding to the probe leading to highest blockade of iodoacetamide-desthiobiotin (IA-DTB) did not exceed 20%, and that the average IA-DTB blockade by the probe was > 66.7% with > 2.5-fold stereoselectivity.

**Cloning, Expression and Purification of recombinant PSME4-20S CP complex for cryoEM**

The cDNA of PSME4 (coding for PSME4) was obtained from in-house human cDNA library (Max Planck Institute of Biochemistry) and was cloned into the baculovirus transfer vector pACEBac1 through Gibbson Assembly. Baculovirus transfer vectors for expression of 20S CPs (Addgene_214138, Addgene_214140, Addgene_214137) have been described previously^3^. The PSME4-c20S complex was co-expressed in Hi5 cells using standard protocols upon which the cells were harvested and resuspended in lysis buffer (25 mM HEPES pH 7.5, 150 mM NaCl, 1mM DTT). Cell lysis was performed by sonication, and the lysate was cleared by centrifugation at 4º C at 50,000g for 1 hour. Subsequently, the cleared lysate was then applied to Strep-Tactin® Sepharose (IBA) for standard affinity purification followed by size exclusion chromatography on Superose6 Increase 10/300 column (Cytiva) equilibrated with a buffer containing 25 mM HEPES, 150 mM NaCl and 0.5 mM TCEP. Individual fractions were analyzed by SDS-PAGE and fractions corresponding to the PSME4–20S CP complex were pooled for further experiments.

**Covalent probing of PSME4-20S CP with PipAc2-4**

Buffer exchange of all steps was carried out utilizing Zeba Microspin Desalting columns. Prior to compound treatment, the purified PSME4-20S complex was incubated with 10 mM DTT and incubated at RT for 10 min. The complex was then buffer exchanged into reaction buffer containing 25 mM HEPES pH 7.5, 150 mM NaCl. Subsequently, free DTT was removed, and a 20-fold molar excess of PipAc2-4 was added, and the reaction was incubated at 30º C for 1 h. Following incubation, the sample was desalted again into the reaction buffer to remove unbound PipAc2-4.

**Structure determination by cryo-EM**

3.5 µl of the PipAc2-4 bound PSME4–20S CP complex at 0.7 mg/ml was applied to glow discharged Quantifoil R1.2/1.3 Cu-200 grids (Quantifoil Micro Tools GmbH) and plunge frozen in liquid ethane/propane mix with Vitrobot Mark IV (Thermo Fisher Scientific) at 4º C and 95% humidity. Grids were screened for optimal conditions via SerialEM v4.1on a TFS Glacios cryo-TEM operated at 200 kV and equipped with a Gatan K3 direct electron detector utilizing SerialEM v4.1. High resolution data collection was performed on a TFS Titan Krios G2 cryo-TEM operated at 300 kV equipped with a Gatan K3 direct electron detector and Bio Quantum post-column energy filter at a pixel size of 0.8222 Å. A total dose of 50 e- per Å2 was fractionated over 40 frames, with a target defocus range of -0.6 and -1.8 µm. The initial processing steps included batch motion correction, contrast transfer function estimation, manual curation of micrographs, blob picking, template picking, 2D classification, ab-initio model generation, and heterogeneous refinement that ultimately provided a particle stack with PSME4-bound 20S CP. To better visualize the PipAc2-4 molecule, a mask was created for the PSME4 density and used to perform local refinement followed by particle subtraction. The subtracted volume for PSME4 along with the classified particle stack was then used for subsequent 3D classification and local refinements to obtain the best density for PipAc2-4 bound to PSME4 within the PSME4-20S CP complex. All processing steps were carried out using CryoSparc (v5.0.6)^4^. For details, see Supplementary Fig. S3.

The final local refined map of PSME4 was used to build the atomic model of PipAc2-4-bound PSME4 using PDB:6KWY as starting reference. The initial model was first fit into the map in ChimeraX and iteratively refined in Coot v1.2^5^ and refined in Phenix v1.2.1^6^.

**Purification of proteasome complexes (MAPP).**

Cells were lysed with 25 mM HEPES, pH 7.4, 10% glycerol, 5 mM MgCl2, 1 mM ATP, and 1:400 protease-inhibitor mixture (Calbiochem), then homogenized through freeze–thaw cycles and passed through a needle. The lysates were cleared by 30-min centrifugation at 21,130g at 4 °C. Lysates were treated with 2 mM 1,10-phenanthroline (Sigma), cross-linked with 0.5 mM DSP (Thermo Fisher Scientific) for 30 min at room temperature, and quenched in 100 mM Tris-HCl, pH 8, 5 mM L-cysteine for 10 min at room temperature. For immunoprecipitation, the lysates were then incubated with Protein G–Sepharose beads (Santa Cruz) with antibodies to PSMA1 and eluted with 100 mM Tris-HCl, pH 8, 8M urea and 50 mM DTT for 30 min at 37 °C. Subsequently, 1% trifluoroacetic acid (TFA) was added. Aliquots of each elution fraction were analyzed by SDS–PAGE to evaluate yield and purity.

**Purification and concentration of proteasome peptides.**

A critical step in our procedure is the separation of peptides from the proteins eluted in the proteasome pulldown. MAPP analyzes endogenously cleaved peptides, whereas the proteasome complex and associated proteins are physically excluded. Immunoprecipitated proteasomes and their encompassed peptides were loaded on C18 cartridges (Waters) that were prewashed with 80% acetonitrile (ACN) in 0.1% TFA, then washed with 0.1% TFA only. After loading, the cartridges were washed with 0.1% TFA. Peptides were eluted with 30% ACN in 0.1% TFA. Protein fractions were eluted with 80% ACN in 0.1% TFA.

**Mass spectrometry sample processing**

Protein fraction after proteasome purification; Proteins were denatured by 8M Urea for 30 min at room temperature, reduced with 5 mM dithiothreitol (Sigma) for 1hr at room temperature and alkylated with 10 mM iodoacetamide (Sigma) in the dark for 45 min at room temperature. Samples were diluted to 2M urea with 50mM ammonium bicarbonate. Proteins were then subjected to digestion with trypsin (Promega; Madison, WI, USA) overnight at 37°C at 50:1 protein:trypsin ratio, followed by a second trypsin digestion for 4 hr. The digestions were stopped by addition of trifluroacetic acid (1% final concentration). Following digestion, peptides were desalted using Oasis HLB, μElution format (Waters, Milford, MA, USA). The samples were vacuum dried and stored in -80˚C until further analysis.

Total proteomics; Lysates in 5% SDS in 50 mM Tris-HCl were incubated at 96 °C for 5 min, followed by six cycles of 30 s of sonication (Bioruptor Pico, Diagenode, USA). Proteins were reduced with 5 mM dithiothreitol and alkylated with 10 mM iodoacetamide in the dark. Each sample was loaded onto S-Trap microcolumns (Protifi, USA) according to the manufacturer’s instructions. In brief, after loading, samples were washed with 90:10% methanol/50 mM ammonium bicarbonate. Samples were then digested with trypsin for 1.5 h at 47 °C. The digested peptides were eluted using 50 mM ammonium bicarbonate; trypsin was added to this fraction and incubated overnight at 37 °C. Two more elutions were made using 0.2% formic acid and 0.2% formic acid in 50% acetonitrile. The three elutions were pooled together and vacuum-centrifuged to dry. Samples were kept at -80 °C until analysis.

**Liquid chromatography mass spectrometry**

Peptide fraction; ULC/MS grade solvents were used for all chromatographic steps. Each sample was loaded using split-less nano-Ultra Performance Liquid Chromatography (10 kpsi nanoAcquity; Waters, Milford, MA, USA). The mobile phase was: A) H2O + 0.1% formic acid and B) acetonitrile + 0.1% formic acid. Desalting of the samples was performed online using a reversed-phase Symmetry C18 trapping column (180 µm internal diameter, 20 mm length, 5 µm particle size; Waters). The peptides were then separated using a T3 HSS nano-column (75 µm internal diameter, 250 mm length, 1.8 µm particle size; Waters) at 0.35 µL/min. Peptides were eluted from the column into the mass spectrometer using the following gradient: 4% to 35%B in 120 min, 35% to 90%B in 5 min, maintained at 90% for 5 min and then back to initial conditions.

The nanoUPLC was coupled online through a nanoESI emitter (10 μm tip; New Objective; Woburn, MA, USA) to a quadrupole orbitrap mass spectrometer (Q Exactive Plus, Thermo Scientific for PSME4 overexpression and Exploris480 for PipAc2-4 treated cells) using a FlexIon nanospray apparatus (Proxeon).

Data was acquired in data dependent acquisition (DDA) mode, using a Top10 method. MS1 resolution was set to 70,000 (at 400m/z), mass range of 375-1650m/z, AGC of 3e6 and maximum injection time was set to 100msec. MS2 resolution was set to 17,500, quadrupole isolation 1.7m/z, AGC of 1e5, dynamic exclusion of 40sec and maximum injection time of 150msec.

**Mass spectrometry data analysis.**

Raw data were analyzed in MaxQuant software (version 1.6.0.16) with the default parameters for the analysis of the proteasomal peptides, except for the following: unspecific enzyme, LFQ minimum ratio count of 1, minimum peptide length for unspecific search of 6, maximum peptide length for unspecific search of 40, and match between runs enabled. A stringent false discovery rate (FDR) of 1% was applied for peptide identification. For the analysis of tryptic digests, the default parameters were set, apart from a minimum peptide length of 6. Masses were searched against the human proteome database from UniprotKB (last update April 2020).

**Proteomics processing and label-free quantification.**

Peptides resulting from MaxQuant were initially filtered to remove reverse sequences and known MS contaminants. For MAPP peptide fraction we removed antibody and proteasome peptides as contaminants. To decrease ambiguity, we allowed peptides that had at least two valid LFQ intensities out of sample replicates, and we included razor peptides, which belong to a unique MaxQuant 'protein group'. MAPP protein intensities were inferred with MaxQuant. For graphical representation, intensities were log transformed, and in Python v3.6, zero intensity was imputed to a random value chosen from a normal distribution of 0.3 s.d. and downshifted 1.8 s.d. The presence of missing values reflects both technical and biological variation in the samples. In cases of matched samples ratios were calculated per pair and paired t tests were used. Otherwise, ratios were calculated based on the median of each group and a non-paired student t test was used.

**Peptide cleavage analysis**

For each peptide, its absence or presence in each sample was annotated (scored 0 or 1) and the C-terminal amino acid was determined (N-terminal amino acid is the amino acid before the peptide start or cleavage site). Cysteines were not quantified as they might be affected by the crosslinker. Per sample, the relative frequency of each amino acid was calculated and standardized on the amino acid level for the heat map representation (*z*-scores). Heat maps were generated with the ComplexHeatmap (v.2.18.0) package with row clustering using Euclidean distances.

**Signatures, statistical analysis and data visualization.**

Statistical analyses were performed in R v 3.6.2 and GraphPad Prism v 7.04. In R data was visualized using the complexheatmap ^7^ and ggplot ^8^ packages. Defaults were used unless otherwise noted. Protein annotation and gene ontology analysis were performed with the gene-set enrichment analysis^9^ using the annotation sets indicated in the figures.

**Antibodies and plasmids**

Rabbit anti PSME4 HPA060922 1:1000 WB (Sigma), Rabbit anti PSME4 ab5620 1:100 IP (Abcam), Mouse anti β Actin ab170325 1:5000 WB (Abcam), Rabbit anti PSMB8/LMP7 13635 1:1000 WB, Rabbit anti 6XHisTag ab137839 1:1000 WB (Abcam), Mouse anti FlagTag F3165 1:1000 WB (Sigma), Rabbit anti STAT1 14994 1:1000 WB (Cell signaling), Mouse anti STAT3 91391:1000 WB 1:100 IP (Cell signaling), Rabbit anti pSTAT3 (Y705) 9145 1:1000 WB (Cell signaling, and Mouse anti GAPDH MAB374 1:1000 WB (Sigma)

Human HLA-A,B,C-PE (W6/32, Biolegend 311406), FC, 1:20; mouse CD45.2-PerCP/Cy5.5 (104, eBioscience 45-0454-82), FC, 1:200; mouse CD3e-Super Bright 436 (145-2C11, eBioscience 62-0031-82), FC, 1:100; mouse CD4-APC/Cy7 (GK1.5, Biolegend 100414), FC, 1:200; mouse CD8a-Super Bright 702 (53-6.7, Biolegend 100748), FC, 1:200; mouse CD279 (PD-1)-PE (RPM1-30, eBioscience 12-9981-81), FC, 1:100; mouse IFNg-APC (XMG1.2, eBioscience 17-7311-82), FC, 1:160; mouse/human CD44-FITC (IM7, Biolegend 103005), FC, 1:200; mouse CD62L-PE/Cy7 (MEL-14, Biolegend 104418), FC, 1:6000; mouse FoxP3-Pe (FJK-16s, eBioscience 12-5773-82), FC, 1:200; mouse H-2Db-FITC (KH95, Biolegend 111505), FC, 1:200; mouse H-2kb-PE (AF6-88.5.5.3, eBioscience 12-5958-82), FC, 1:500; Zombie Aqua (Biolegend 423102), FC, 1:300; Propidium Iodide (Sigma Aldrich P4170), FC, 1 ug/mL; CFSE (Biolegend 423801), FC, 5 uM. Mouse anti alpha-6 (PSMA1), produced from hybridoma, was a kind gift from Keiji Tanaka (1:1000 WB). IgG1k isotype antibody (BioLegend, BLG-400139) was used as a control of MAPP method. More information regarding antibodies can be found in supplementary table S6.

Goat anti Mouse 488 A11029 1:400, Goat anti mouse 647 A31571 1:400 (Invitrogen), Goat anti Rabbit 647 AB150075 1:400 (Abcam), Goat anti mouse HRP 115-035-205 1:5000, Goat anti Rabbit HRP 111-035-003 1:5000 (Jackson labs).

MISSION shRNA targeting mouse or human PSME4 or RFP were obtained from Sigma (TRCN0000176569, TRCN0000178428, TRCN0000158223, TRCN0000157073).

pcDNA3.1_PSME4 ^10^ was transfected into A549/KP1.9 cells with Lipofectamine 2000 (Thermo Fisher). ORF of PSME4 was cloned to pcDNA3.1(Zeo+) with 6XHisTag at the N- terminus of the protein. Site directed mutagenesis method was preformed to create C1001A point mutation (GenScript). Both plasmids were validated by sequencing.

pmIL-6promoterEGFP was a gift from Nicolai van Oers (Addgene plasmid # 112896; http://n2t.net/addgene:112896; RRID: Addgene_112896)^11^

**Cell culture and drug treatments**

A549 were originally obtained from ATCC grown in DMEM and used as model systems for cell biology studies including western blot, biochemistry, and imaging. KP1.9, kindly provided by Alfred Zippelius, grew in Iscove’s MDM. Cells were routinely tested for mycolplasma contamination. Media were supplemented with 10% fetal bovine serum, 1% Penicillin/streptomycin and L-glutamine (2mmol/l) (Biological industries), unless otherwise indicated, at 37 °C with 5% CO2.

For TNF-α/IFN-γ activation cells were allowed to seed overnight, and subsequently treated with TNF-α (PeproTech, human; 300-01A or mouse; 315-01A) and IFN-γ (PeproTech, human; 300-02 or mouse; 315-05) at 20 ng/ml and 10 ng/ml, respectively, for 24h.

For PipAc’s compounds cells were allowed to seed overnight and subsequently treated with 10 µM for 18h unless stated differently. For proteasome inhibition we used bortezomib 100 nM for 18h.

Cell death assessment was done by trypan blue staining and counted with Countess II™ automated cell counter (Thermo Fisher) and by Cell titerGlo assay (Promega). Proliferation of mammalian cells were measured by KIT8 assay (Sigma).

**PSME4-20S and 20S complexes purification**

20S and PSME4-20S proteasomes were purified from the testes of C57BL/6JOlaHsd mice bearing a chromosomal insertion encoding a C-terminal FLAG tag in PSMB2 (β4). Proteasome complexes were purified according to a previously described protocol^12^. Briefly, samples were subjected to two sequential rounds of DEAE anion-exchange chromatography followed by size-exclusion chromatography. PSME4-20S complexes were further purified by Mono Q chromatography. PSME4-containing fractions were monitored by immunoblotting with anti-PSME4 antibodies. The composition and integrity of the purified complexes were verified by SDS-PAGE with Coomassie blue staining, and immunoblotting (Extended Data Fig. 3). Recombinant 20S human proteasomes were purchased (Enzo, BML-PW8720) and used as a reference.

**Proteasome cleavage reporter assay**

Cells were lysed with 25 mM HEPES, pH 7.4, 10% glycerol, 5 mM MgCl2, 1 mM ATP, and 1:400 protease-inhibitor mixture (Calbiochem), then homogenized through freeze–thaw cycles and passed through a needle. The lysates were cleared by 30-min centrifugation at 21,130g at 4 °C to remove cell debris. Protein concentration of the supernatant was determined by a NanoDrop spectrophotometer (Thermo Fisher Scientific) by measuring the absorbance at 280 nm.

Proteasomal activity in cell fractions was determined by cleavage of the fluorogenic precursor substrate Suc-Leu-Leu-Val-Tyr-AMC (Suc-LLVY-AMC), Ac-Pro-Ala-Leu-AMC (Ac-PAL-AMC), Z-Leu-Leu-Glu-AMC (Z-LLE-AMC), Ac-Arg-Leu-Arg-AMC (Ac-RLR-AMC), Ac-Ala-Asn-Trp-AMC (Ac-ANW-AMC) (Boston Biochem), Z-Leu-Leu-Glu-βNA (Z-LLE- βNA), Ac-Nle-Pro-Nle-Asp-AMC (Ac-nLPnLD-AMC) (Bachem). 10 μM substrate was added to 10 µg total protein/well, and incubated in a reaction buffer (50 mM Hepes, pH 7.5, 1 mM DTT, 5 mM MgCl2 and 2 Mm ATP). Fluorescent increase resulting from degradation of peptide-AMC at 37°C was monitored over time by means of a fluorometer (Synergy H1 Hybrid Multi-Mode Microplate Reader, BioTek) at 340 nm excitation and at 460 nm emission, using the proteasome inhibitor, MG132 (Calbiochem) as background. Resulting product curves were followed for up to 3.5 hours. Each value of fluorescence intensity represents a mean value obtained from three independent experiments.

**Activity assay of isolated proteasomes**

Black 96-wells ELISA plate was coated with mouse anti alpha-6 (PSMA1) antibody in 50mM carbonate-bicarbonate pH 9.5 Buffer, overnight. The following day cells were lysed with 25 mM HEPES, pH 7.4, 10% glycerol, 5 mM MgCl_2_, 1 mM ATP and 1:400 protease-inhibitor mixture (Calbiochem), homogenized through freeze–thaw cycles and passed through a needle (25G). The lysates were cleared by 30-min centrifugation at 21,130*g* at 4 °C. Protein concentration was assessed using a BC Assay Protein Quantitation Kit (Interchim). Black 96-wells coated plate was washed 3 times with PBS contains 0.05% Tween-20 and blocked with 3%BSA in PBS 1h in RT. 20 μg cell lysate was added to the black 96-well plate and incubated overnight on an orbital shaker. The next day the plate was washed three times in PBS contained 0.05% Tween-20 and one time with PBS. Proteasome activity was determined as described above, by cleavage of the fluorogenic precursor substrates in a reaction buffer (50 mM HEPES, pH 7.5, 1 mM dithiothreitol, 5 mM MgCl_2_ and 2 mM ATP). The increase in fluorescence at 37 °C was monitored over time using a fluorometer (Synergy H1 Hybrid Multi-Mode Microplate Reader, BioTek) at 340 nm excitation and 460 nm emission. The resulting product curves were followed for up to 3.5 h. Each fluorescence intensity represents a mean value obtained from three or more independent experiments.

**IL-6 release assay (ELISA)**

IL-6 secretion was measured by ELISA assay (ELISA MAX™ Deluxe Set Human IL-6, Biolegend) by manufacture procedure. Plates were scanned using the Synergy H1 Hybrid Multi-Mode Microplate Reader (BioTek) at 450 wavelength absorbance.

**Immuno-blotting**

Cells were washed twice with PBS and lysed in STET buffer (50 mM Tris-HCl, pH 87.5, 150 mM NaCl, 2 Mm EDTA 1% Triton-X and 1:400 protease-inhibitor mixture (Calbiochem)). Protein concentration was assessed using BC Assay Protein Quantitation Kit (Interchim). 20 μg of total protein was separated by SDS–PAGE on 4-20% gradient ExpressPlus PAGE M42015 (GenScript) and transferred onto PVDF membranes using iBlot™ 3 Gel Transfer Device (Thermo Fisher Scientific). The membranes were blocked in 5% milk prepared in TBS-0.1% Tween and incubated in primary antibodies overnight at 4°C followed by washing and incubation with secondary antibody. Blots were developed using the ChemiDoc XRS+ Imaging System (Bio-Rad) and band intensities were quantified with imageJ analyzer software.

**Immuno-blotting for phosphorylated proteins**

Cells were washed twice with PBS, and lysed in Cellytic™ M, Cell Lysis Reagent, Suitable for mammalian cell lysis and protein solubilization (Sigma-Aldrich), 1:10 Phosphatase inhibitor (PhosSTOP, Sigma-Aldrich), and 1:400 Protease Inhibitor Cocktail III (Sigma-Aldrich). Next, cells were incubated for 15 minutes on ice, followed by addition of Benzonase nuclease (1:1000) and incubation for 10 minutes at 30 degrees. Lysates were cleared by centrifugation (16,000g, 10 min, 4°C) to remove cell debris. Protein concentration was assessed using BC Assay Protein Quantitation Kit (Interchim). 20 μg of total protein was separated by SDS–PAGE on 4-20% gradient ExpressPlus PAGE M42015 (GenScript) and transferred onto Nitrocellulose membranes using iBlot™ 3 Gel Transfer Device (Thermo Fisher Scientific). The membranes were blocked in 3% BSA prepared in TBS-0.1% Tween and incubated in primary antibodies overnight at 4°C followed by washing and incubation with secondary antibody. Blots were developed using the ChemiDoc XRS+ Imaging System (Bio-Rad) and band intensities were quantified with imageJ analyzer software.

**Immuno-precipitation**

Cells lysed in 25 mM HEPES, pH 7.4, 10% glycerol, 5 mM MgCl2, 1 mM ATP, and 1:400 protease-inhibitor mixture (Calbiochem), then homogenized through freeze–thaw cycles and passed through a needle. The lysates were cleared by 30-min centrifugation at 21,130g at 4 °C to remove cell debris. Pre-cleared cell lysates were prepared by incubating with Protein A/G conjugated agarose beads for 30 min at 4°C with slow rotation. 500 µg pre-cleared cell lysates were then added to 5 µL of fresh Protein A/G conjugated agarose beads along with 4 µg of antibodies (as described). The mixture was incubated over-night at 4°C with slow rotation followed by three 1 ml washes with chilled PBS buffer. The target conjugates were then eluted from beads by mixing protein loading dye (with reducing agents) and heating for 5 min at 95°C followed by denaturing Western blot.

**Non-denaturing immuno-precipitation**

The protocol was adapted from^14^, in brief: Cells were washed twice with PBS, and lysed in Cellytic™ M, Cell Lysis Reagent, Suitable for mammalian cell lysis and protein solubilization (Sigma-Aldrich), 1:10 Phosphatase inhibitor (PhosSTOP, Sigma-Aldrich), and 1:400 Protease Inhibitor Cocktail III (Sigma-Aldrich). Next, cells were incubated on ice for 15 minutes, followed by the addition of Benzonase nuclease (1:1000) and incubation for 10 minutes at 30 degrees. Lysates were cleared by centrifugation (16,000g, 10 min, 4°C) to remove cell debris. Protein concentration was assessed using a BC Assay Protein Quantitation Kit (Interchim). 500 μg of cleared lysates were then incubated with 4 μg of antibody on ice. After 1h protein A/G MagBeads (GenScript) were added and then incubated overnight at 4°C with head to tail rotation, followed by three 1 ml washes with a chilled TBS buffer. The target conjugates were then eluted from beads by mixing 4X LDS Sample Buffer (GenScript) (with reducing agents 30mg/ml DTT and denaturant 4M Urea) and heating for 2 min at 95 °C followed by denaturing western blot.

**qPCR analysis**

RNA was extracted using Direct-zol™ RNA MiniPrep R2051 (ZYMO research) mRNA levels were ascertained by RT using High-Capacity cDNA Reverse Transcription Kit (ThermoFisher) real time quantitative PCR using sybr-green (Kapa Biosystems) and the StepOnePlus™ Real-Time PCR Systems (Life Technologies) using primers listed in supplementary table S6.

All values were normalized to the mRNA abundance of housekeeping genes (RPS18 in Human; GAPDH in Mouse). Each primers pair was calibrated using the Absolute Quantification program with increasing concentrations of cDNA.

**Peptide stripping by acid wash**

To remove HLA-bound peptides from the cell surface, cells were washed twice with PBS and then incubated for 1 min with citrate phosphate buffer (260 mM citric acid and 123 mM disodium phosphate (pH 3)), at room temperature. Afterwards, cells were washed twice with their relevant medium, supplemented with 10% FBS, 2 mM glutamine, and 1 mM sodium pyruvate. Subsequently, cells were resuspended in fresh medium and plated in 6 wells plate.

**Surface HLA staining in flow cytometry**

A549 and KP1.9 cells were treated with hTNFα/hIFNγ (as described above) or mTNFα/mIFNγ (as described above), respectively, for 24 hr. Cells were harvested and stained with Live/Dead Aqua staining kit from BioLegend as per manufacturer’s protocol. Approximately 1x10^6^ cells were washed, blocked with 2% FCS for 5 min and stained with anti-HLA-A/B/C-PE (A549) or anti-MHCI-Kb-PE/MHCI-Db-FITC (KP1.9) in 100ul for 30 min. Cells were washed and acquired.

**Orthotopic lung cancer model and Immunotherapy**

Mouse experiments were conducted according to approved experimental procedures (Approval numbers 04400520-2 and 04990620-1). Male mice (C57BL/6JRccHsd, age 8-10 weeks) were injected with 0.25x10^6 KP1.9 i.v., either KP1.9/shCtrl or KP1.9/shPSME4. Mice were monitored three times a week for general wellbeing, signs of sickness and body weight changes. Antibody treatments started on day 8 after tumor injection at a dose of 200ug/mouse i.p., twice a week. Mice were removed from the study once their body weight dropped by 20% of the highest weight.

Anti-IL6 antibody was from Bio X Cell, cat BE0046, clone MP5-20F3, lot 670721A1. Rat IgG1 isotype control was from Bio X Cell, cat BE0088, clone HRPN, lot 724921M1. Anti-PD-1 antibody was from ichor bio, cat ICH1132, clone RPM1-14, lot 0423L615. Rat IgG2a isotype control was from ihor bio, cat ICH2244, clone 1-1, lot 0623L605. Antibodies were diluted fresh in PBS before each injection.

**Sucutaneous cancer model (priming and boosting)**

Mouse experiments were conducted according to approved experimental protocol (Approval number 00850126-1). Male C57Bl/6JRccHsd mice, aged 8-10 weeks, were injected with 0.25x10^5 KP1.9/shCtrl cells to prime the immune system to the cancer. After 3 weeks, KP1.9/shCtrl cells were pre-treated for 24 hours with compound PIPAc2-4 at 10 µM concentration or DMSO for control. KP1.9/shPSME4 also received DMSO at the same concentration. Cells were harvested, washed extensively, resuspended in PBS and injected to mice bilateral to the flanks, each site receiving 3x10^6 cells. After 48 hours, tumors were harvested and their weights noted. Of each pair, one tumor went for processing in western blot analysis, the other for flow cytometric analysis. For this, tumors were digested with a mix of 200U/mL Collagenase 4 (Worthington Biochemical LS004188), 100 ug/mL DNase I (Sigma-Aldrich, DN25), in empty RPMI1640 medium at 37C on a shaking platform for 30 minutes. Samples were strained and stained for flow cytometry. After surface staining, cells were fixed and permeabilized for FoxP3 staining using the Ebioscience Foxp3 / Transcription Factor Staining Buffer Set (00-5523-00) according to the manufacturer protocol. Cells were finally washed and acquired on a Attune Nxt flow cytometer.

**Survival plots**

Survival data were analyzed using the Kaplan–Meier method and compared between treatment groups using the log-rank (Mantel–Cox) test. Survival time was defined as the number of days from tumor implantation until death or censoring (alive at study endpoint). For multiple treatment comparisons, pairwise survival analyses were performed using the pairwise_survdiff function from the *survival* R package with rho = 0 (log-rank test). Adjusted *p*-values from pairwise comparisons were symbolically coded according to significance thresholds (**** < 0.0001, *** < 0.001, ** < 0.01, * < 0.05, + < 0.1, – ≥ 0.1) for visualization.

**Software**: R Version 4.5.1; RStudio 2025.09.1, 401; gridExtra_2.3; survival_3.8-3; survminer_0.5.1; ggpubr_0.6.1; lubridate_1.9.4; forcats_1.0.0; stringr_1.5.2; dplyr_1.1.4; purrr_1.1.0; readr_2.1.5; tibble_3.3.0; ggplot2_3.5.2; tidyverse_2.0.0; tidyr_1.3.1; readxl_1.4.5

**Tissue processing and flow cytometry staining**

Peripheral blood was washed and red blood cells removed by ACK lysis buffer (150 mM NH4Cl, 10 mM KHCO3, 0.1 mM EDTA in 0.1x PBS-/-) and washed in flow cytometry buffer (PBS-/-, 0.5% BSA, 2mM EDTA).

Spleens and lungs were weighed before further processing. The left lung lobe was used for histological analysis of tumor development (hematoxylin and eosin) after fixation in 4% formaldehyde (Biolabs, Israel). Right lung lobes were minced and digested with 5 mL digestion buffer: Collagenase 4 (200U/mL, Worthington Biochemicals), DNase I (100 ug/mL, Sigma Aldrich) in PBS++ supplemented with 2 mM CaCl2 for 20 minutes at 37°C under shaking. Single cell suspensions of spleen and lungs were obtained by straining cells through 100 um strainers and washed in cold flow cytometry buffer. Red blood cells were removed by ACK lysis buffer, washed with flow cytometry buffer and strained again.

For intracellular cytokine and transcription factor stainings, cells were incubated for 4 hours with Brefeldin A (Biolegend) and Monensin (Sigma Aldrich). First, viability staining was done with Zombie Aqua (Biolegend) according to the protocols. Cell surface staining was performed in flow cytometry buffer in 100 μL at appropriate dilutions. When needed, cells were fixed and permeabilized with the FoxP3 transcription factor kit from eBioscience (Thermo Fisher Scientific) and stained for FoxP3 and IFNg according to the manufacturers protocol. Samples were acquired on an Attune Nxt with Autosampler and analyzed in FlowJo (V10.7.1, Becton, Dickinson and Company). Exemplary gating strategies are provided Figures S10, S20 and S22)

**Cytokine and chemokine secretion**

Tumor microenvironment; 10,000 KP1.9 WT cells were plated onto a 96-well plate and allowed to attach overnight. The following day, splenocytes were isolated from naïve mice, or from mice bearing KP1.9-WT, -PSME4 KD, or -PSME4 OE tumors. Red blood cells were lysed using ACK buffer, cells were washed in PBS, and 100,000 splenocytes were plated on top of the tumor cells in RPMI medium + beta-mercaptoethanol (50uM). Co-culture was maintained for 48h or 7 days. The medium was collected at 48h and 7days by centrifugation. Cytokines and chemokines concentration in the tumor microenvironment were determined by 36-Plex Mouse ProcartaPlex (ThermoFisher, EPX360–26092–901) as per manufacturer’s instructions. Samples were acquired on MAGPIX System (Luminex) and analysed by MILLIPLEX Analyst V5.1 Flex.

Tumor cells secretion; 10,000 KP1.9 shCtrl or KP1.9 PSME4 KD cells were plated onto a 96-well plate and allowed to attach overnight. The following day cells were treated with mTNFα and mIFNγ mix for 24h in the last 6h ONX-0914 (1µM) (APExBIO, A4011) was added to desired wells. Cells were washed twice and incubated for 24h in fresh medium. The medium was collected flowed be centrifugation. Cytokines and chemokines secretion were determined by 36-Plex Mouse ProcartaPlex (ThermoFisher, EPX360–26092–901) as per manufacturer’s instructions. Samples were acquired on MAGPIX System (Luminex) and analyzed by MILLIPLEX Analyst V5.1 Flex.

**MILLIPLEX MAP cell signaling portfolio**

MILLIPLEX MAP Multi path Phospho 9 plex (48-680MAG; Millipore) and MILLIPLEX MAP Multi path Total 9 plex (48-681MAG; Millipore) assays were performed. All samples were run in duplicates on a single plate per panel. Kits were purchased from EMD Millipore Corporation and used according to the manufacturer’s recommendations. Briefly, samples were lysed in kit’s buffer and mixed with antibody-linked magnetic beads in a 96-well plate and incubated overnight at 4°C with shaking. Cold and room temperature incubation steps were performed on an orbital shaker at 500 rpm. Plates were washed twice with a wash buffer. Following a 1-hour incubation at room temperature with biotinylated detection antibody, streptavidin-PE was added for 30 minutes with shaking. Plates were washed as above, and reading buffer was added to wells for reading with a lower bound of 50 beads per sample per protein. Samples were acquired on MAGPIX System (Luminex) and analyzed by MILLIPLEX Analyst V5.1 Flex.

**Statistics and reproducibility**

All experiments were performed in three individual replicates unless otherwise mentioned. For each experiment, all compared conditions were analyzed by MS simultaneously. For the NSCLC study, nine samples from tumors and adjacent tissue were analyzed. The samples were processed independently and analyzed by MS simultaneously to maintain comparability across samples and minimize batch effects.

**Data availability**

The mass spectrometry proteomics data related to cysteine-directed ABPP has been deposited to the ProteomeXchange Consortium via the PRIDE^13^ partner repository with the dataset identifier PXD069990.

Cryo-EM Maps and Model have been deposited to the EMDB and PDB for release upon publication (validation reports are provided to reviewers). To assist with review, these are available at: <https://datashare.biochem.mpg.de/s/66zwTppkJC5oJRk>

Mass spectrometry data (listed in the Supplementary Tables MS datasets) have been deposited with the ProteomeXchange Consortium via the PRIDE^13^ partner repository, with the dataset identifier PXD083978 (<https://www.ebi.ac.uk/pride/>). Previously published data for clinical samples proteasome composition partner repository with the primary accession code [PXD019573](http://proteomecentral.proteomexchange.org/cgi/GetDataset?ID=PXD019573)  (<https://www.ebi.ac.uk/pride/>).
